# SUMOylation of PTEN promotes DNA end resection through directly dephosphorylating 53BP1 in homologous recombination repair

**DOI:** 10.1101/2023.02.06.527258

**Authors:** Jianfeng He, Yanmin Guo, Rong Deng, Lian Li, Caihu Huang, Ran Chen, Yanli Wang, Jian Huang, Jinke Cheng, Guo-Qiang Chen, Junke Zheng, Xian Zhao, Jianxiu Yu

## Abstract

Homologous recombination (HR) repair for DNA double-strand breaks (DSBs) is critical for maintaining genome stability and cell survival. Nuclear PTEN plays a key role in HR repair, but the underlying mechanism remains largely elusive. We find that SUMOylated PTEN promotes HR repair but represses non-homologous end joining (NHEJ) repair by directly dephosphorylating 53BP1. During DNA damage responses (DDR), p14ARF was phosphorylated and then interacted efficiently with PTEN, thus promoting PTEN SUMOylation as an atypical SUMO E3 ligase. Interestingly, SUMOylated PTEN was subsequently recruited to the chromatin at DNA-break sites. This was because that SUMO1 conjugated to PTEN was recognized and bound by the SUMO-interacting motif (SIM) of BRCA1, which has been located to the core of 53BP1 foci on the chromatin during S/G2 stage. Further, these chromatin-loaded PTEN directly and specifically dephosphorylated pT543 of 53BP1, resulting in the dissociation of 53BP1-complex, which facilitated DNA end resection and ongoing HR repair. The SUMOylation-deficient PTEN^K254R^ mice also showed decreased DNA damage repair *in vivo.* Blocking PTEN SUMOylation pathway with either an SUMOylation inhibitor or a p14ARF(2-13) peptide sensitized tumor cells to chemotherapy. Our study therefore provides the new mechanistic understanding of PTEN in HR repair and clinical intervention of chemo-resistant tumors.

## Introduction

DNA lesions caused by environmental or endogenous genotoxic insults are major threats to genomic integrity(***Roos et al., 2016, Tubbs et al., 2017***). DNA double strand breaks (DSBs) are the most deleterious DNA lesions, which cause gene mutation, cell death, development disorder and tumor predisposition if not repaired correctly and promptly(***Chen et al., 2020, Lord et al., 2016***). There are two major pathways for DSB repair, homologous recombination (HR) and non-homologous end joining (NHEJ) repair(***Scully et al., 2019***). 53BP1, a pro-choice of DSBs, promotes NHEJ repair through inhibiting recruitment of HR repair factors including BRCA1 and CtIP to DSB sites in G1 stage(***Panier et al., 2014***). DNA damage-induced phosphorylation at multiple sites in the N terminal of 53BP1 mediates its interaction with downstream factors RIF1 and PTIP(***Callen et al., 2013***). RIF1 recruits the Shieldin complex, of which the subunit SHLD2 directly binds ssDNA and blocks DNA end resection, and loss of this complex dramatically increases HR repair(***Gupta et al., 2018, Noordermeer et al., 2018, Zimmermann et al., 2013***). On the other hand, HR repair depends on the exist of sister chromatid, which occurs mainly in S/G2 stage. BRCA1, a critical regulator of HR repair, promotes multiple steps including DNA end resection, RAD51 loading and ssDNA strand pairing(***Chapman et al., 2012, Huen et al., 2010***). BRCA1 can recruit a ubiquitin E3 ligase UHRF1 to mediate polyubiquitination of RIF1, resulting in RIF1 dissociation from 53BP1 and thus promoting HR repair in S/G2 stage(***Zhang et al., 2016***). Moreover, BRCA1 can also facilitate dephosphorylation of 53BP1 during S/G2 stage(***Isono et al., 2017***). The region coded by *exon11* of *BRCA1* is required for the dephosphorylation of 53BP1 and RIF1 release from DNA breaks, however the underlying molecular mechanism remains largely elusive(***Nacson et al., 2020, Nacson et al., 2018***).

PTEN, a dual phosphatase, is frequently deleted, mutated or downregulated in a variety of human tumors(***Lee et al., 2018***). In cytoplasm, PTEN antagonizes PI3K-AKT signaling through its lipid phosphatase activity, while loss of which markedly promotes tumor cell proliferation(***Lee et al., 2018***). It has been well documented that the nuclear PTEN plays a critical role in maintaining the genome stability, centrosome stability, replication stress recovery and DSB repairs(***Bassi et al., 2013, He et al., 2015, Ho et al., 2020, Ma et al., 2019, Shen et al., 2007, Wang et al., 2015, Zhang et al., 2019***). Post-translational modifications (PTMs) of PTEN including SUMOylation, phosphorylation and methylation are involved in DNA damage and repair(***Bassi et al., 2013, Huang et al., 2012, Ma et al., 2019, Zhang et al., 2019***). SUMOylation has been extensively studied in DNA damage repair(***Dhingra et al., 2019, Schwertman et al., 2016***), and many DNA damage response (DDR) associated proteins including CtIP, BMI1, BLM, RAD52 and TOP2A can be induced occurring SUMOylation by replication stress, DSB and DNA crosslinking(***Choi et al., 2013, Cremona et al., 2012, Ismail et al., 2012, Ouyang et al., 2009, Soria-Bretones et al., 2017, Tian et al., 2021***). Especially, PTEN SUMOylation plays a key role in repairing for DSB, but the underlying mechanism remains unexplored. In addition, there still remains dispute about the function of PTEN protein phosphatase activity in DDR.

Here we provided evidences that SUMOylation of PTEN was increased by SUMO-E3-like p14ARF in DDR. SUMOylated PTEN was recognized and then recruited by the N-terminal SUMO-interacting motif (SIM) of BRCA1 to the chromatin. PTEN located at chromatin directly and specifically dephosphorylated pT543 of 53BP1, leading to RIF1 release and therefore facilitating DNA end resection. PTEN^K254R^ knock-in mice model also showed HR deficiency in DDR *in vivo*. Notably, inhibiting SUMOylation of PTEN by either a SUMOylation inhibitor or a peptide p14ARF(2-13) sensitized tumor cells to DNA damage agents, which might provide a new therapeutic strategy for clinical intervention of chemo-resistant tumor cells.

## Results

### PTEN promotes HR repair through facilitating DNA end resection

We first validated the role of PTEN in DSB repair. As shown in (***Figure 1—figure supplement 1A-C***), knockdown of PTEN in DU145 cells indeed delayed DSB repair after the treatment with Zeocin (a radiomimetic reagent), as measured by ionizing radiation-induced foci (IRIF) of γH2AX and 53BP1(***Bassi et al., 2013, Zhang et al., 2019***). Either PTEN knockdown in DU145 and HeLa cells or PTEN knockout in MEFs (Mouse embryonic fibroblasts) dramatically reduced the numbers of RAD51 (a key HR repair regulator) foci (***Figure 1—figure supplement 1D-F***), which was consistent with previous reports(***Bassi et al., 2013, Ma et al., 2019***). In addition to RAD51 filament formation, DNA end resection, a key step prior to RAD51, is also critical for the choice of DSB repair pathway and can be regulated in many ways(***Symington, 2016***). To test whether PTEN regulates DNA end resection, we detected the phosphorylation level of RPA32(S4/8), which is a marker of DNA end resection in DSB repair(***Nimonkar et al., 2011***), to show that knockdown of PTEN dramatically inhibited pS4/8-RPA32 in HeLa cells after irradiation (IR) treatment (***Figure 1A***), suggesting that PTEN is involved in the regulation of DNA end resection in DSB repair.

**Figure 1.**
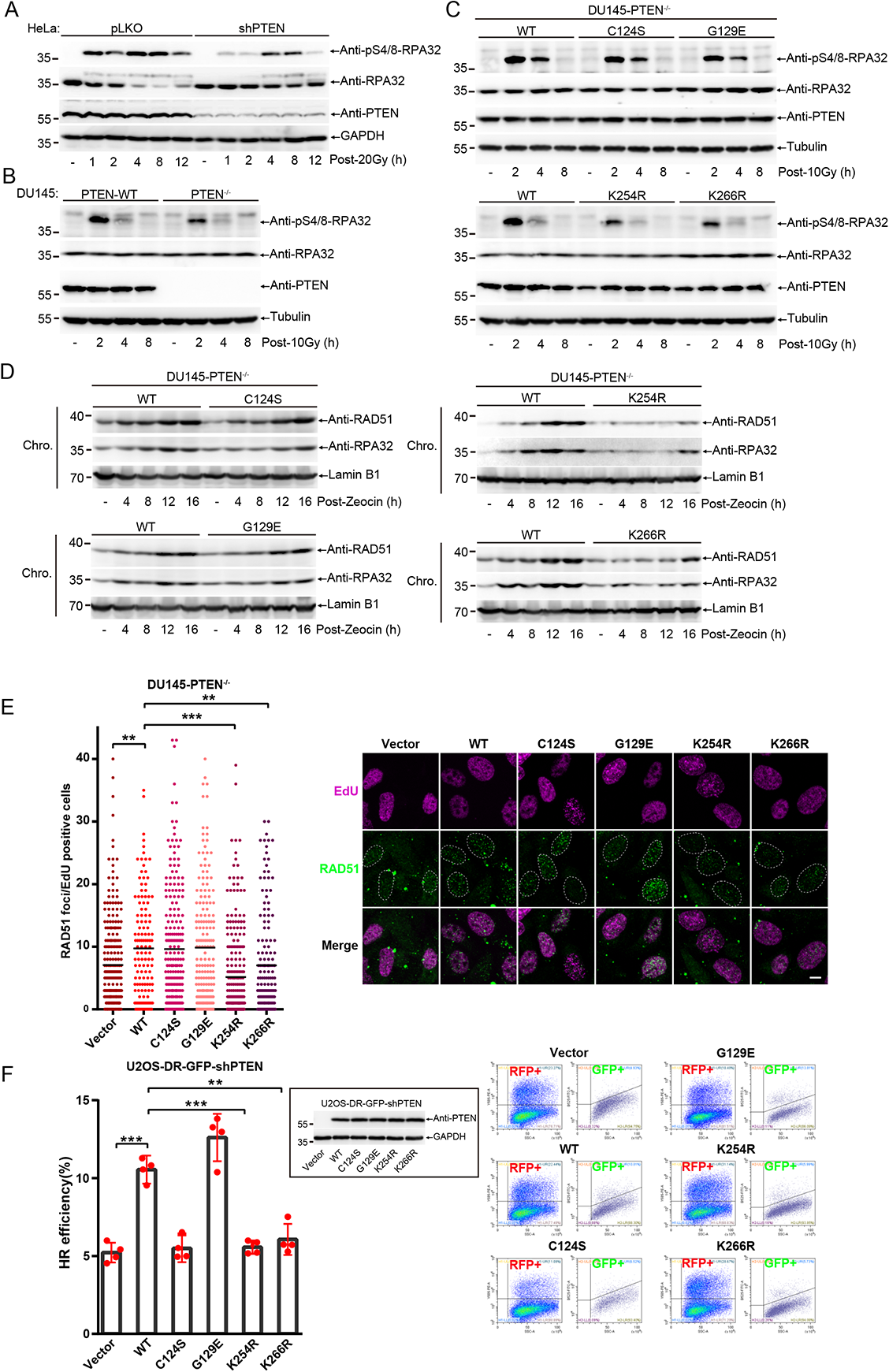
PTEN promotes HR repair through facilitating DNA end resection. (**A**) Immunoblot of pS4/8-RPA32 of HeLa-pLKO and -shPTEN cells treated with 20 Gy and recovery for indicated time. (**B**) Immunoblot of pS4/8-RPA32 of DU145-PTEN^WT^ and -PTEN^−/−^ cells treated with 10 Gy and recovered for indicated time. (**C**) Immunoblot of pS4/8-RPA32 of DU145-PTEN^−/−^ cells stably re-expressed PTEN^WT^, ^K254R^ and ^K266R^ cells after treated with 10 Gy. (**D**) Immunoblot of chromatin associated RPA32 and RAD51 in DU145 cells after treated with Zeocin (200 μg/mL) for 1 h and recovered for indicated time. (**E**) RAD51 foci were quantified in DU145 cells treated with 5 Gy and recovery for 6 h and then presented with dot graph (n(Vector)=223, n(WT)=135, n(C124S)=221, n(G129E)=187, n(K254R)=244, n(K266R)=167). Representative images of RAD51 foci were shown at right panel. scale bar, 20 μm. (**F**) HR efficiency were detected in U2OS-DR-GFP-shPTEN cells stably re-expressing PTEN^WT^, PTEN^C124S^, PTEN^G129E^, PTEN^K254R^ and PTEN^K266R^ (n=4 for each group). Inset: immunoblot of PTEN in U2OS-DR-GFP-shPTEN cells stably re-expressing indicated PTEN mutants. Left panel: quantification of HR efficiency shown as bar graph. Right panel: representative images of FACS. Unpaired Student’s t-test was used (**p< 0.01, ***p< 0.001) and data were shown as mean or mean±s.d. **Figure 1-source data 1.** Western blots of Figure 1.

To further identify whether both the phosphatase activity and SUMOylation of PTEN are required for its role in DNA end resection, we used the CRISPR/Cas9 system to knockout PTEN in DU145 cells and then stably re-expressed PTEN^WT^, PTEN^C124S^ (dual phosphatase dead), PTEN^G129E^ (lipid phosphatase dead, but protein phosphatase still active), PTEN^K254R^ (SUMO-site mutant) and PTEN^K266R^ (SUMO-site mutant) (***Figure 1—figure supplement 1G***). As expectedly, knockout of PTEN decreased IR-induced pS4/8-RPA32 in DU145 cells (***Figure 1B***). There were little differences on pS4/8-RPA32 among PTEN^WT^, PTEN^C124S^ and PTEN^G129E^ after IR treatment, however pS4/8-RPA32 was clearly decreased in PTEN^K254R^ and PTEN^K266R^ compared to that in PTEN^WT^ (***Figure 1C***). Similar results of PTEN^K254R^ and PTEN^K266R^ in decreasing pS4/8-RPA32 were also observed by using other DNA damage reagents including Zeocin and Camptothecin (CPT) in DU145 (***Figure 1—figure supplement 1H and I***), and IR in PC3 cells (***Figure 1—figure supplement 1J***). Thus, above results suggest that SUMOylation but not phosphatase activity of PTEN is essential for DNA end resection.

Determination of chromatin associated proteins has been often applied to monitor DNA damage repair process, for examples, chromatin loading of RPA32 and RAD51 can represent HR efficiency (***Andegeko et al., 2001, Ma et al., 2019, Mendez et al., 2000, Yang et al., 2017***). When compared with PTEN^WT^, mutants PTEN^K254R^ and PTEN^K266R^ greatly inhibited the chromatin loading of RPA32 and RAD51 in DU145 cells induced by Zeocin (***Figure 1D***), Etoposide (***Figure 1—figure supplement 1K***) and CPT (***Figure 1—figure supplement 1L***), whereas PTEN^C124S^ and PTEN^G129E^ seemed not affect (***Figure 1D***). Furthermore, we also investigated IRIF of RAD51 to show the similar pattern of results that numbers of RAD51 foci were comparable among PTEN^C124S^, PTEN^G129E^ and PTEN^WT^, but decreased in PTEN^K254R^ and PTEN^K266R^ (***Figure 1E***). These results suggest that SUMOylation but not phosphatase activity of PTEN affects the chromatin loading of RPA32 and RAD51.

In addition to RAD51 foci as a marker of HR repair, an HR reporter of DR-GFP was employed to detect the overall efficiency of HR repair (***Xie et al., 2018***). In accordance with previous reports, HR efficiency was reduced after PTEN knockdown (***Figure 1—figure supplement 1M***). Surprisingly, HR efficiency was also compromised in PTEN^C124S^, as like PTEN^K254R^ and PTEN^K266R^, but not in PTEN^G129E^, suggesting PTEN protein phosphatase was indispensable for it function in HR repair efficiency despite its little influence on chromatin loading of RPA32 and RAD51 (***Figure 1F***). Taken together, these data provide substantial evidences that PTEN promotes HR repair partially through enhancing DNA end resection, which is dramatically abolished when its SUMO-sites mutated.

### DNA damage promotes PTEN chromatin loading by inducing its SUMOylation

As previous reported (***Cremona et al., 2012***), DNA damage stimuli can strongly induce SUMOylation of proteins involved in different DNA damage repair pathways. To further investigate the induction and turnover of PTEN SUMO modification in DNA damage repair, we overexpressed His-SUMO1 and Flag-PTEN in 293T cells. After Zeocin treatment, cells were collected and lysed at different recovery time as indicated, and His-SUMO1 modified proteins were enriched with Ni^2+^-NTA agarose beads (***Figure 2A***) and Co-IP (***Figure 2—figure supplement 1A***) methods under denatured condition. Interestingly, SUMOylated PTEN was significantly increased overtime by Zeocin treatment. Similarly, CPT treatment also enhanced SUMOylation of PTEN (***Figure 2B***). These data strongly demonstrated that DNA damage stimuli promoted PTEN SUMOylation. To further determine which SUMO-site is responsible for SUMO1 conjugation induced by DNA damage and whether its phosphatase activity is involved in this process, we overexpressed His-SUMO1 and Flag-tagged PTEN^C124S^, PTEN^G129E^, PTEN^K254R^ or PTEN^K266R^ in 293T cells and detected SUMOylated PTEN with the method of Ni^2+^-NTA agarose pull down. In contrast to mutations C124S and G129E, both mutations of K254R and K266R led to significant reduction of PTEN SUMOylation induced by Zeocin, suggesting that these two SUMO-sites were critical for DDR (***Figure 2C***).

**Figure 2.**
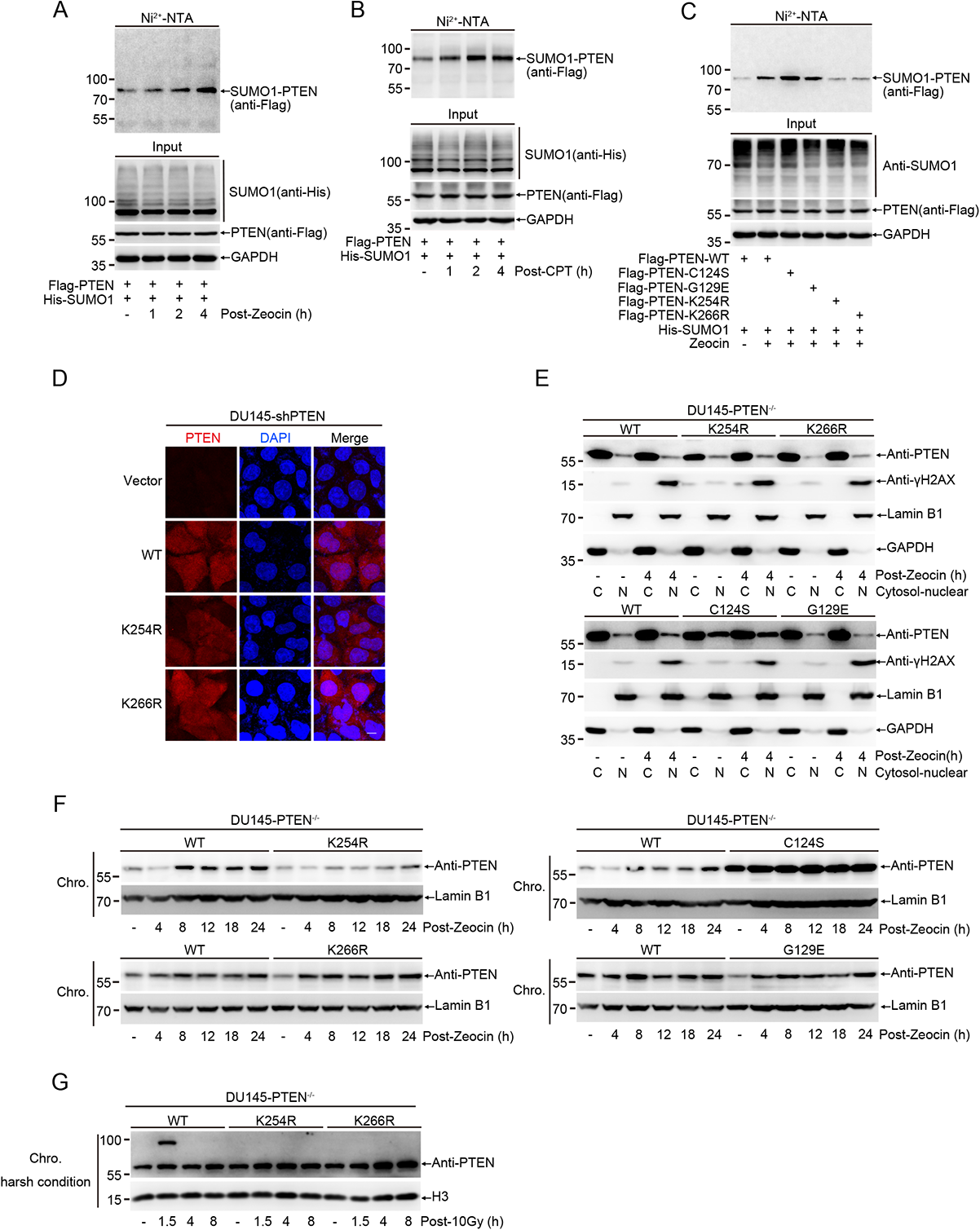
DNA damage promotes PTEN chromatin loading by inducing its SUMOylation. (**A**) 293T cells transfected with His-SUMO1 and Flag-PTEN were treated with Zeocin (200 μg/mL) for 1h and recovered for indicated time. His-SUMO1 conjugates were pulled down with Ni^2+^-NTA beads and SUMOylated PTEN were analyzed with immunoblot. (**B**) 293T cells transfected with His-SUMO1 and Flag-PTEN were treated with CPT (20 μM) for 1h and recovered for indicated time. His-SUMO1 conjugates were pulled down with Ni^2+^-NTA beads and SUMOylated PTEN were analyzed with immunoblot. (**C**) 293T cells transfected with His-SUMO1 and Flag-PTEN^WT^, ^C124S^, ^G129E^, ^K254R^, ^K266R^ were treated with Zeocin (200 μg/mL) for 1h and recovered for 4 h, Ni^2+^-NTA pulldown were used to analyze PTEN SUMOylation. (**D**) Images of staining of PTEN in DU145-shPTEN cells stably re-expressed PTEN-WT, K254R and K266R. (**E**) Nuclear-Cytosol separation was performed in DU145-PTEN^−/−^ cells stably re-expressing PTEN^WT^, ^C124S^, ^G129E^, ^K254R^ and ^K266R^ treated with Zeocin (400 μg/mL) for 1 h and recovery for 4 h. Localization of PTEN was detected with immunoblot. (**F**) Immunoblot of chromatin associated PTEN in DU145 cells treated with Zeocin (400 μg/mL) for 1 h and recovery for indicated time. (**G**) Chromatin protein separation were performed at harsh condition in DU145-PTEN^−/−^ cells stably re-expressing PTEN^WT^, ^K254R^ and ^K266R^ after treatment with 10 Gy. PTEN tightly associated with chromatin was detected with immunoblot. **Figure 2-source data 1.** Western blots of Figure 2.

One previous study reported that K254R mutation prevented PTEN nuclear localization (***Bassi et al., 2013***), however our results of immunofluorescence showed no changes in the localization between PTEN^WT^ and PTEN^K254R^ or PTEN^K266R^ in DU145 cells under normal condition (***Figure 2D***). Moreover, the results of nuclear-cytosol separations also revealed that the localizations of PTEN were almost not affected in PTEN-WT and mutants including PTEN^K254R^, PTEN^K266R^, PTEN^C124S^ and PTEN^G129E^ even after treatments with Zeocin (***Figure 2E***) and CPT (***Figure 2—figure supplement 1B***). More interestingly, the nuclear location of PTEN^C124S^ was markedly higher than that of all others (***Figure 2E, Figure 2—figure supplement 1B***), the underlying mechanism should be further studied.

Although SUMO-site mutations K254R and K266R had little influence on PTEN nuclear localization, we wanted to assess whether SUMOylation of PTEN influences its chromatin loading under DNA damage. First, we observed that PTEN was indeed recruited into the DNA-damage location induced by the laser micro-irradiation (***Figure 2—figure supplement 1C***). Second, Zeocin (***Figure 2F***) and CPT (***Figure 2—figure supplement 1D***) treatments induced PTEN accumulation on the chromatin. The chromatin loading of PTEN^K254R^ was significantly suppressed whereas that of PTEN^G129E^ and PTEN^K266R^ was similar with PTEN^WT^ after Zeocin and CPT treatments. For the case of PTEN^K266R^, it was unexpectedly and might be a different mechanism from PTEN^K254R^. In accordance with enhanced nuclear localization of PTEN-C124S, the chromatin loading of PTEN^C124S^ was remarkably increased (***Figure 2F***), but how the mutation C124S to increase the nuclear localization and chromatin loading of PTEN was not clear. Third, to further validate whether SUMOylated PTEN can directly accumulate on the chromatin, we separated chromatin associated proteins under harsh condition. After IR treatment, one shifted band with higher molecular weight than normal PTEN was clearly observed in re-expression of PTEN^WT^ but not SUMO-site mutants PTEN^K254R^ and PTEN^K266R^ in DU145-PTEN^−/−^ cells (***Figure 2G***). Given that endogenous *PTEN* gene in DU145 cells was knocked out with CRSPR/Cas9 technique, this shift band from re-expression of normal PTEN^WT^ was most likely to be SUMOylated-PTEN other than variants of PTEN such as PTEN α or β isoform.

Collectively, all above results demonstrate that DNA damage promotes PTEN SUMOylation and especially K254-SUMOylation of PTEN is essential for its chromatin loading.

### p14ARF is a novel SUMO E3 ligase to mediate PTEN SUMOylation during DDR

p14ARF, a well-known tumor suppressor, is an atypical SUMO E3 ligase for promoting SUMOylation of its binding proteins such as MDM2, NPM and EGR1 (***Ozenne et al., 2010, Tago et al., 2005, Yu et al., 2009***). Interestingly, we found that PTEN SUMOylation was dramatically enhanced by overexpression of p14ARF (***Figure 3A, Figure 3—figure supplement 1A***) whereas suppressed by knockdown of p14ARF (***Figure 3B***). Since p14ARF promotes SUMOylation of target proteins *via* direct interaction, we assumed p14ARF might interact with PTEN. Reciprocal co-IP and GST-pull down results showed that p14ARF directly bound to PTEN in cells and *in vitro* (***Figure 3C and D***, ***Figure 3—figure supplement 1B***). To identify the interaction region between p14ARF and PTEN, we generated a series of truncates (***Figure 3—figure supplement 1C***) and performed co-IP. The results showed that two major regions 2-14aa and 82-101aa in p14ARF were required for their interaction. Deletion of either one reduced their interaction and deletion of both completely abolish their interaction (***Figure 3E***, ***Figure 3—figure supplement 1D* and *E***). Moreover, the ability of p14ARF to enhance PTEN SUMOylation was indeed weakened when deletion of either one region. Although both regions contributed to PTEN SUMOylation, p14ARF(△2-14) seemed to be more effective than p14ARF(△82-101) in suppression of PTEN SUMOylation (***Figure 3F***). In addition, we identified that the C-terminal region 188-403aa of PTEN mediated its interaction with p14ARF (***Figure 3—figure supplement 1F***).

**Figure 3.**
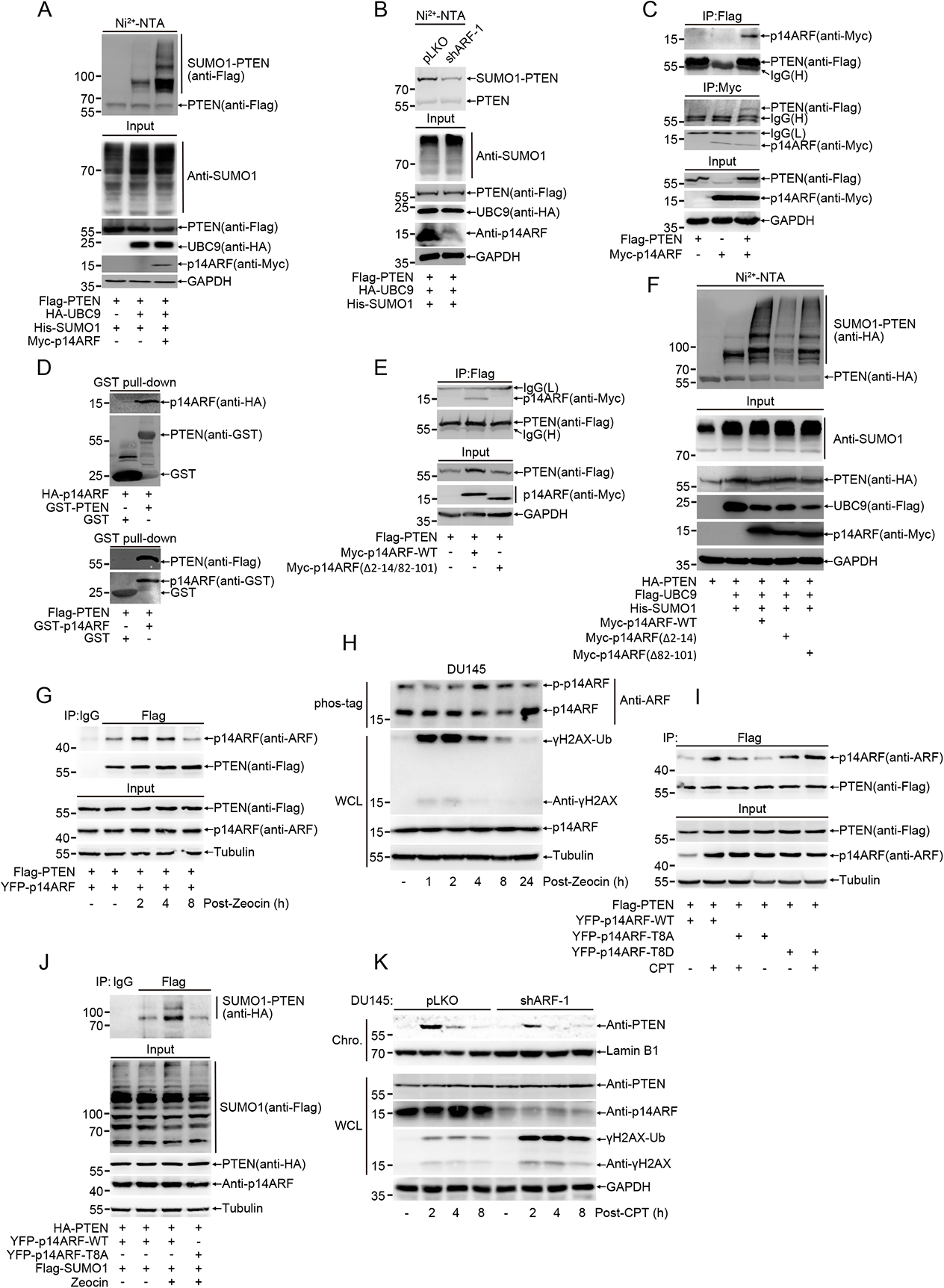
p14ARF is a novel SUMO E3 ligase to mediate PTEN SUMOylation during DDR. (**A**) 293T cells were transfected with His-SUMO1, HA-UBC9, Myc-p14ARF and Flag-PTEN for 48 h. His-SUMO1 conjugates were pulled down with Ni^2+^-NTA beads and SUMOylated PTEN were analyzed with immunoblot. (**B**) 293T-pLKO or shARF-1 cells were transfected with His-SUMO1, HA-UBC9 and Flag-PTEN for 48 h. His-SUMO1 conjugates were pulled down with Ni^2+^-NTA beads and SUMOylated PTEN were analyzed with immunoblot. (**C**) 293T cells were transfected with Myc-p14ARF and Flag-PTEN for 48 h. Co-IP were used to detect interaction between PTEN and p14ARF. (**D**) GST pull-down were used to detect interaction between PTEN and p14ARF. Upper panel: GST-PTEN was purified from BL21 and incubated with 293T lysis which transfected with HA-p14ARF for 48 h. Lower panel: GST-p14ARF was purified from BL21 and incubated with 293T lysis which transfected with Flag-PTEN for 48 h. (**E**) Truncated Myc-p14ARF and Flag-PTEN were transfected into 293T cells, interaction domain between PTEN and p14ARF were identified with Co-IP. (**F**) Truncated Myc-p14ARF, His-SUMO1, Flag-UBC9 and HA-PTEN were transfected into 293T cells. His-SUMO1 conjugates were pulled down with Ni^2+^-NTA beads and SUMOylated PTEN were analyzed with immunoblot. (**G**) Interaction between PTEN and p14ARF post CPT (20 μM) treatment were detected with Co-IP in 293T cells. (**H**) Phosphorylation of p14ARF after Zeocin (400 μg/mL) treatment for 1 h and recovery for indicated time was detected with phos-tag gel. (**I**) Interaction of PTEN and p14ARF-WT, T8A and T8D were detected under normal condition or CPT (20 μM) treatment with Co-IP in 293T cells. (**J**) 293T cells were transfected with Flag-SUMO1, Myc-p14ARF-WT or T8A and HA-PTEN for 48 h. Flag-SUMO1 conjugates were immunoprecipitated and SUMOylated PTEN were analyzed with immunoblot. (**K**) Chromatin loading of PTEN were detected with immunoblot in DU145-pLKO and shARF-1 cells after treatment with CPT (20 μM) for 1 h and recovery for indicated time. **Figure 3-source data 1.** Western blots of Figure 3.

Since DNA damage stimuli can induce PTEN SUMOylation, we wondered whether p14ARF is involved in this process. In response to treatments with both Zeocin (***Figure 3G***) and CPT (***Figure 3—figure supplement 1G***), the interaction of PTEN and p14ARF was obviously enhanced. Given that phosphorylation signal is critical for DDR and there is exactly one threonine (T8) located in p14ARF(2-14aa), which can be phosphorylated (***Fontana et al., 2018***), so we detected the phosphorylation of p14ARF with the method of Phos-tag gel and showed a clear shifted band of p14ARF, which was increased after Zeocin treatment (***Figure 3H***), suggesting that DNA damage induced phosphorylation of p14ARF. The mutation T8A at p14ARF significantly inhibited the interaction between PTEN and p14ARF, on the contrary, the mutation T8D enhanced their interaction not only under normal condition but also after treatment with CPT (***Figure 3I***). The mutation T8A of p14ARF also attenuated its ability to promote PTEN SUMOylation after Zeocin treatment (***Figure 3J***). Consistently, knockdown of p14ARF inhibited PTEN chromatin loading while overexpression of p14ARF increased its loading (***Figure 3K, Figure 3—figure supplement 1H-J***). Knockdown of p14ARF also sensitized DU145 cell to Cisplatin (***Figure 3—figure supplement 1K***). All these results suggest that p14ARF is a functional SUMO E3 ligase responsible for promoting SUMOylation of PTEN during DDR.

### PTEN relieves HR repair barrier posted by 53BP1 through directly dephosphorylating pT543-53BP1

Given that HR repair efficiency is compromised when PTEN lacking protein phosphatase activity (***Bassi et al., 2013***) (***Figure 1F***), next we tried to identify potential protein substrates targeted by PTEN in DNA damage repair. 53BP1, a key negative regulator of HR repair, is phosphorylated at multiple S/TQ sites which are essential for recruitment of downstream effectors (***Callen et al., 2013***). Thus, we first examined the phosphorylation levels of 53BP1 in cells after treatment with CPT or Zeocin. The pT543-53BP1 level was dramatically increased in PTEN-knockdown DU145 and HeLa cells (***Figure 4A, left panels***) whereas significantly decreased in PTEN overexpression in PC3 cells which are devoid of endogenous PTEN (***Figure 4A, right and upper panel***) after treatment with CPT (for 30 min). Moreover, we also detected the pT543-53BP1 level in PTEN-knockout DU145 cells after treatment with higher concentration of CPT for 1 h, to show that the pT543-53BP1 level was higher and lasted longer in PTEN-knockout cells than that of PTEN-WT cells (***Figure 4A, right and lower panel***). Consistent with above results, when treated with Zeocin, the pT543-53BP1 level was increased in PTEN-knockdown DU145 and HeLa cells (***Figure 4—figure supplement 1A***), while weakened in PTEN overexpression in PC3 cells (***Figure 4—figure supplement 1B***). In addition, we confirm that the intensity of pT543-53BP1 foci was stronger in PTEN-knockdown DU145 than control cells ***Figure 4B***, ***Figure 4—figure supplement 1C***) by staining with antibody pT543-53BP1.

**Figure 4.**
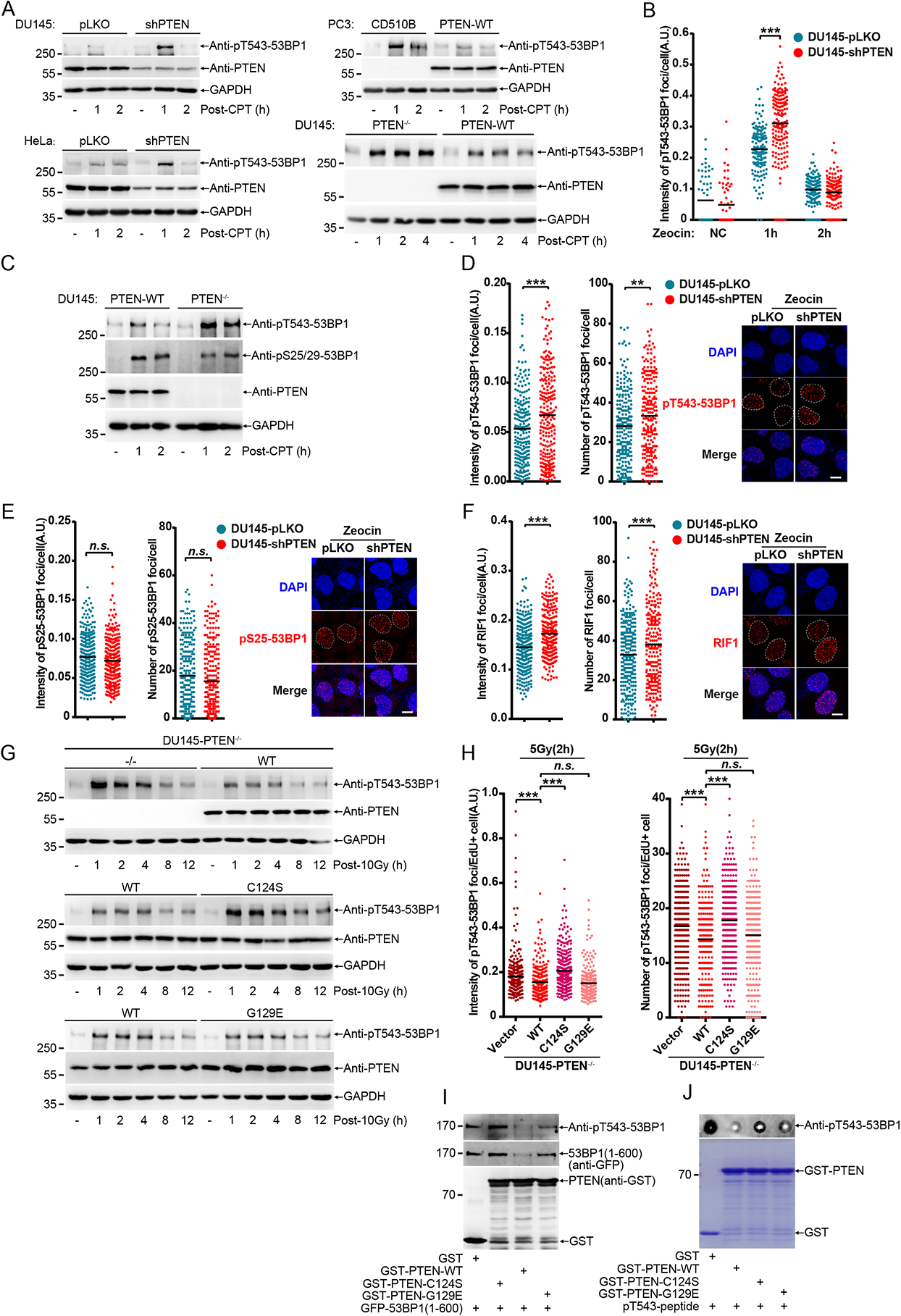
PTEN relieves HR barrier posted by 53BP1 through directly dephosphorylating pT543-53BP1. (**A**) Immunoblot of pT543-53BP1 in DU145 (top left), HeLa (lower left) cells and PC3 (top right) after treatment with CPT (2 μM) for 30 min and recovery for indicated time. pT543-53BP1 was also detected in DU145 (lower right) cells after treated with CPT (20 μM) for 1 h and recovery for indicated time. (**B**) Dot graph of pT543-53BP1 foci intensity in DU145 cells after Zeocin (100 μg/mL) treatment for 30 min and recovery for indicated time (DU145-pLKO(n(NC)=49, n(1h)=152, n(2h)=158), DU145-shPTEN(n(NC)=61, n(1h)=173, n(2h)=168)). (**C**) Immunoblot of pT543-53BP1 and pS25/29-53BP1 in DU145 cells after treatment with CPT (20 μM) for 1 h and recovery for indicated time. (**D**) Dot graph of pT543-53BP1 foci number and intensity after Zeocin (200 μg/mL) treatment for 1 h and recovery for 2 h (DU145-pLKO(n=228), DU145-shPTEN(n=229)). Representative images were shown at right panel. (**E**) Dot graph of pS25-53BP1 foci number and intensity after Zeocin (200 μg/mL) treatment for 1 h and recovery for 2 h (DU145-pLKO(n=267), DU145-shPTEN(n=292)). Representative images were shown at right panel. (**F**) Dot graph of RIF1 foci number and intensity after Zeocin (200 μg/mL) treatment for 1 h and recovery for 2 h (DU145-pLKO(n=273), DU145-shPTEN(n=239)). Representative images were shown at right panel. (**G**) Immunoblot analysis of pT543-53BP1 level in DU145-PTEN^−/−^ cells stably re-expressed PTEN-WT, C124S and G129E after 10 Gy treatment and recovery for indicated time. (**H**) Dot graph of pT543-53BP1 foci number and intensity in DU145-PTEN^−/−^ cells stably re-expressed PTEN-WT, C124S and G129E after 5 Gy treatment and recovery for 2 h (n(Vector)=327, n(WT)=222, n(C124S)=322, n(G129E)=269). (**I**) in vitro phosphatase assay. GFP tagged phosphorylated 53BP1(1-600) were purified from 293T cells. GST-PTEN^WT^, ^C124S^ and ^G129E^ were purified from E.coli BL21. (**J**) in vitro phosphatase assay was performed with synthesized pT543-53BP1 polypeptides. Unpaired Student’s t-test was used (**p< 0.01, ***p< 0.001) and data were shown as mean. n.s.: not significant. **Figure 4-source data 1.** Western blots of Figure 4.

Since it has been reported that PTEN loss led to the increase of pS25/29-53BP1 after treatment with etoposide and the mutation S25A enhanced RIF1 recruitment during DNA damage repair (***Callen et al., 2020, Zhang et al., 2019***), thus we also detected the dynamics of pS25/29-53BP1 in PTEN-knockdown or -knockout cells after treatment with CPT or Zeocin. Indeed, pS25/29-53BP1 was induced in both PTEN-WT and PTEN-knockout DU145 cells after treatment with CPT. Increased pS25/29-53BP1 level was the same and even a little down whereas as expectedly, increased pT543-53BP1 level was higher in PTEN-knockout DU145 when compared with PTEN-WT DU145 (***Figure 4C***). Furthermore, after treatment with Zeocin, the number and intensity of pT543-53BP1 foci were expectedly increased in DU145-shPTEN cells compared to those in DU145-pLKO cells (***Figure 4D***); in contrast, there was little difference in the number and intensity of pS25-53BP1 foci between DU145-pLKO and DU145-shPTEN cells (***Figure 4E***). These results demonstrate that PTEN regulates the level of pT543-53BP1 other than pS25/29-53BP1 in DDR.

As pT543-53BP1 helps to recruit RIF1 to DNA breaks (***Callen et al., 2013***), so we stained RIF1 in DU145-pLKO and shPTEN cells after IR treatment. In accordance with the pT543-53BP1 levels, the number and intensity of RIF1 foci were both increased (***Figure 4F***). It has been well documented that the 53BP1-RIF1-shieldin axis forms a barrier to inhibit HR repair through suppressing DNA end resection and HR repair mainly occurs at S/G2 of cell cycle (***Feng et al., 2013, Mirman et al., 2018, Noordermeer et al., 2018, Zimmermann et al., 2013***), we wondered that whether PTEN in regulation of pT543-53BP1 is dependent on cell cycle. DU145 cells were firstly synchronized at G1/S with double thymidine block, and then directly released 5 hours to enter S phase or synchronized at G1 with lovastatin, respectively. Cyclin A2 was used to show synchronization efficiency (***Figure 4—figure supplement 1D***). The pT543-53BP1 levels were enhanced in both G1 and S phases after treatment with Zeocin (***Figure 4—figure supplement 1E and F***). However, the increased pT543-53BP1 levels in S phase were much weaker in PTEN^WT^ cells than those PTEN^−/−^ cells (***Figure 4—figure supplement 1E***). In contrast, the increased pT543-53BP1 levels in G1 phase were slightly weaker in PTEN^WT^ cells when compared with PTEN^−/−^ cells (***Figure 4—figure supplement 1F***). These data suggest that PTEN regulating pT543-53BP1 is a cell-cycle dependent manner and mainly occurs in S phase.

To further verify if PTEN phosphatase is responsible for dephosphorylating pT543-53BP1, we determined the pT543-53BP1 levels in PTEN^−/−^, PTEN^WT^, PTEN^C124S^ and PTEN^G129E^ DU145 cells. After treatment with IR or Zeocin, the pT543-53BP1 levels in PTEN^−/−^ cells were higher than those in PTEN^WT^ cells, as was expected (***Figure 4G, Figure 4—figure supplement 1G, top panels***). We found that pT543-53BP1 was increased in PTEN^C124S^ but not in PTEN^G129E^ cells when compared with PTEN^WT^ cells (***Figure 4G, Figure 4—figure supplement 1G, top panels, middle and low panels***). As similar to immunoblot results, the number and intensity of pT543-53BP1 foci were also increased in PTEN^C124S^ but not in PTEN^G129E^ cells compared to those in PTEN^WT^ DU145 cells at 2 h after treatment with IR (5 Gy) (***Figure 4H, Figure 4—figure supplement 1H***). Above results indicated that PTEN protein phosphatase was required for dephosphorylation of pT543-53BP1. Thus, we further speculated whether PTEN directly dephosphorylates pT543-53BP1.

To verify above hypothesis, we first purified phosphorylated full-length Flag-53BP1 from 293T cells for the *in vitro* reaction with GST-PTEN^WT^ or GST-PTEN^G129R^ (a dual phosphatase deficient mutant). The following Western blotting results showed that PTEN-WT but not PTEN-G129R efficiently dephosphorylated pT543-53BP1 (***Figure 4—figure supplement 1I***), suggesting that the PTEN phosphatase activity is required for dephosphorylation of pT543-53BP1. Next, we transformed the two truncated forms into 293T cells, and found that GFP-53BP1(1-600) was easily detected by antibody anti-pT543-53BP1 and more strongly after CPT treatment; on the contrary, GFP-53BP1(1-300) could not be detected (***Figure 4—figure supplement 1J***). In order to further distinguish protein phosphatase and lipid phosphatase of PTEN in dephosphorylation of pT543-53BP1, similarly, we purified phosphorylated GFP-53BP1(1-600) from 293T cells for the *in vitro* reaction with GST-PTEN^WT^, GST-PTEN^C124S^ or GST-PTEN^G129E^. Consistent with cellular results, PTEN^C124S^ lost the ability to dephosphorylate pT543-53BP1, whereas PTEN^G129E^ could moderately dephosphorylate pT543-53BP1 but was less efficient than PTEN-WT, which might be because the G129E mutation not only abolished lipid phosphatase of PTEN, but also reduced its protein phosphatase by about a half (***Chia et al., 2010***) (***Figure 4I***). We noticed that GFP-53BP1(1-600) (also Flag-53BP1 in ***Figure 4—figure supplement 1I***) partly degraded after the *in vitro* reaction with GST-PTEN^WT^ and GST-PTEN^G129E^, but not GST-PTEN^C124S^ and GST-PTEN^G129R^, suggesting that phosphorylation modification might be important to maintain 53BP1 protein stability *in vitro.* Moreover, we synthesized a small pT543-peptide (^536^IDEDGEN^543^T(p)QIEDTEP^550^) for the similar *in vitro* reaction with GST-PTEN^WT^, GST-PTEN^C124S^ or GST-PTEN^G129E^, and confirmed that PTEN^WT^ could efficiently dephosphorylate pT543-peptide and PTEN-G129E partly did; on the contrary, PTEN^C124S^ seemed to lose this ability (***Figure 4J***). Thus, above results suggest that PTEN directly dephosphorylates pT543-53BP1 *in vitro* and in DDR.

Taken together, our data demonstrate that PTEN directly dephosphorylates pT543-53BP1 in response to DNA damage, which relieves HR repair barrier posted by 53BP1.

### PTEN chromatin loading is mediated by BRCA1 recruiting SUMOylated PTEN via its N-terminal SIM

BRCA1 can remove HR repair barrier posted by 53BP1 *via* several molecular mechanisms, including suppression of 53BP1 phosphorylation induced by DNA damage (***Isono et al., 2017***), so we wondered whether PTEN is a downstream effector of BRCA1. As expectedly, knockdown of BRCA1 by siRNA strongly increased the pT543-53BP1 level in DU145 after Zeocin treatment (***Figure 5A***). Consistent with this, knockdown of BRCA1 by either siRNA or shRNA displayed the same results of enhancing pT543-53BP1 in U2OS cells after treatment with Zeocin and Etoposide (***Figure 5—figure supplement 1A***). For the other hand, we questioned whether BRCA1 is involved in PTEN chromatin loading. Indeed, chromatin separation experiments showed that knockdown of BRCA1 by shRNA or siRNA significantly suppressed PTEN chromatin loading as well as pS4/8-RPA32 (as a positive control) in DU145 cells after treatment with CPT, indicating that BRCA1 was needed for PTEN chromatin loading during DDR (***Figure 5B, Figure 5—figure supplement 1B***). Therefore, we next tested whether PTEN interacts with BRCA1 and this can be enhanced by SUMOylation. We transfected Flag-PTEN and Myc-BRCA1 with SUMO1 into *Senp^−/−^*-293T cells for 48 h, and then treated with or without CPT. Co-IP/Western blotting results showed that PTEN interacted with BRCA1, and the interaction was moderately enhanced by CPT treatment. Most strikingly, the interaction was strongest when co-transfected with SUMO1 and UBC9 (SUMO-conjugating enzyme E2), which was not increased any more even with CPT treatment (***Figure 5C***). To identify whether the interaction is specifically enhanced by SUMO1 modification, we also detected the ability of other SUMO isoform SUMO2 whose amino acid sequence is a little different from SUMO1. The interaction was much weaker in SUMO2 transfected than that in SUMO1 transfected cells under CPT treatment. Surprisingly, co-transfected with SUMO2 and UBC9 did not enhance the interaction at all (***Figure 5—figure supplement 1C***). These data suggest that the interaction between PTEN and BRCA1 is specifically promoted by SUMO1 modification.

**Figure 5.**
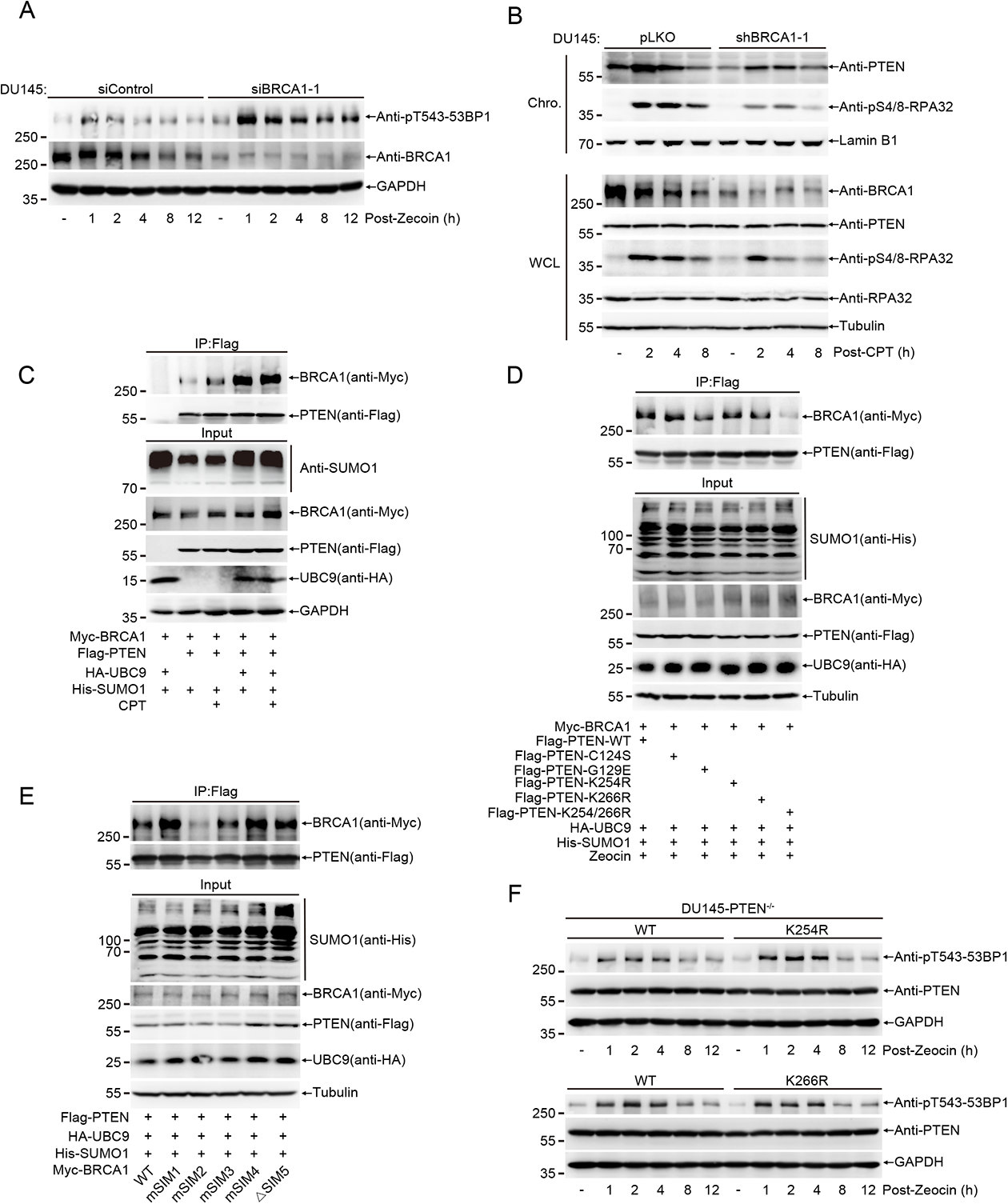
PTEN chromatin loading is mediated by BRCA1 recruiting SUMOylated PTEN via its N-terminal SIM. (**A**) Immunoblot of pT543-53BP1 in DU145 cells in which BRCA1 were knockdown with siControl or siBRCA1-1 after Zeocin (400μg/mL) treatment for 1 h and recovery for indicated time. (**B**) Immunoblot of chromatin loaded PTEN and pS4/8-RPA32 in BRCA1 knockdown DU145 cells after treatment with CPT (20 μM) for 1 h and recovery for indicated time. (**C**) 293T^senp−/−^ cells were transfected with Myc-BRCA1, HA-UBC9, His-SUMO1 and Flag-PTEN for 48 h. Co-IP were performed to identify interaction between PTEN and BRCA1. (**D**) 293T^senp−/−^ cells were transfected with Myc-BRCA1, HA-UBC9, His-SUMO1 and Flag-PTEN (WT or mutants) for 48 h and treated with Zeocin. Co-IP were performed to identify interaction between PTEN (WT or mutants) and BRCA1. (**E**) 293T^senp−/−^ cells were transfected with Myc-BRCA1(WT or SIM mutants), HA-UBC9, His-SUMO1 and Flag-PTEN for 48 h. Co-IP were performed to identify interaction between PTEN and BRCA1 (WT or SIM mutants). (**F**) Immunoblot of pT543-53BP1 in DU145-PTEN^−/−^ cells stably re-expressed PTEN-WT, K254R and K266R after treatment with Zeocin (400 μg/mL) for 1 h and recovery for indicated time. **Figure 5-source data 1.** Western blots of Figure 5.

Given that the SUMO-site mutation of PTEN inhibited its chromatin loading induced by DNA damage, we wondered whether the interaction between PTEN and BRCA1 is directly mediated by SUMO-SIM (SUMO interacting motif), which is an important mechanism to mediate the protein-protein interaction. Indeed, the double mutations K254/266R of PTEN remarkably repressed the interaction although the single mutation K254R or K266R did not inhibit, suggesting SUMO1 modification of PTEN was important for its interaction with BRCA1. Additionally, the lack of lipid phosphatase activity of mutants PTEN^C124S^ and PTEN^G129E^ did not affect the interaction (***Figure 5D***).

To identify SIMs of BRCA1 which are responsible for the interaction with SUMO1 attached to PTEN, we used two software GPS-SUMO and JASSA (***Beauclair et al., 2015, Zhao et al., 2014***) to predict possible SIMs of BRCA1 (***Figure 5—figure supplement 1D and E***). Since depletion of *exon11* of BRCA1 abolishes its suppression on 53BP1 phosphorylation and DNA end resection (***Nacson et al., 2020, Nacson et al., 2018***), we mainly focused on SIMs located in this region, which are marked in red (***Figure 5—figure supplement 1D and E***). Further to find out which SIM is essential for the interaction, we mutated amino acids of SIM1, 2, 3, 4, into alanine, referred as mSIMn, and deleted SIM5-1 and SIM5-2, referred as △SIM5, respectively (***Figure 5—figure supplement 1F***). Co-IP/Western blotting results showed that the interaction was efficiently weakened by mSIM2 (^412^VLDVL^416^--AAAAA) but not by other mutants of BRCA1 (***Figure 5E***).

Since the interaction between PTEN and BRCA1 was dependent on SUMO-SIM, we wondered that the SUMO-site mutant PTEN is also be defective in dephosphorylation of 53BP1. The pT543-53BP1 level was relatively higher in PTEN^K254R^ cells than that in PTEN^WT^ DU145 cells after treatment with Zeocin. However, the pT543-53BP1 level in PTEN^K266R^ cells was comparable with that in PTEN^WT^ cells (***Figure 5F***). These suggest that chromatin loading of PTEN mediated by K254-SUMO1 modification is also very important for its role in dephosphorylation of pT543-53BP1. All above results demonstrate that BRCA1 recruits SUMOylated PTEN to chromatin *via* its N-terminal SIM, thereby dephosphorylating pT543-53BP1 in DDR.

### HR repair is impaired by SUMO-deficient PTEN *in vivo*

To verify whether SUMO-deficient PTEN impairs HR repair *in vivo*, we generated knock-in mice with a point-mutation of *Pten*^K254R^ or *Pten*^K266R^ by using CRISPR-Cas9 technique. *Pten*^K254R^ mice developed normally and was indistinguishable to *Pten*^WT^ mice, but interestingly, *Pten*^K266R^ mice exhibited low birth rate and some of them show abnormal localization of seminal vesicle. MEFs isolated from *Pten*^WT^ and *Pten*^K254R^ mice were verified by DNA sequencing and used to examine DNA damage repair efficiency (***Figure 6—figure supplement 1A***). Compared to *Pten*^WT^ MEFs, *Pten*^K254R^ MEFs showed significant increase of γH2AX and 53BP1 foci (***Figure 6A and B***, ***Figure 6—figure supplement 1B and C***) but decrease of RPA32 and RAD51 foci (***Figure 6C and D***, ***Figure 6—figure supplement 1D and E***) after treatment with IR, indicating *Pten*^K254R^ MEFs were deficient in HR repair. The levels of γH2AX were also higher in *Pten*^K254R^ MEFs compared to those in *Pten*^WT^ MEFs in the late stage after CPT or Zeocin (***Figure 6—figure supplement 1F***). Furthermore, the pS4/8-RPA32 levels were much weaker in *Pten*^K254R^ MEFs than those in *Pten*^WT^ MEFs after treatment with IR of 20 Gy or 30 Gy (***Figure 6E***, ***Figure 6—figure supplement 1G***), which proved that PTEN^K254R^ function in regulation of DNA end resection during HR repair was indeed compromised. Consistent with results of tumor cell lines, PTEN chromatin loading induced by IR and CPT was also inhibited in *Pten*^K254R^ MEFs (***Figure 6F***, ***Figure 6—figure supplement 1H***). Chromatin-bound RPA32 was also decreased in *Pten*^K254R^ MEFs in response to IR treatment, indicating reduced DNA end resection efficiency (***Figure 6F***). Collectively, these data confirm that SUMO-deficient PTEN^K254R^ impairs HR repair by decreasing its chromatin loading and DNA end resection.

**Figure 6.**
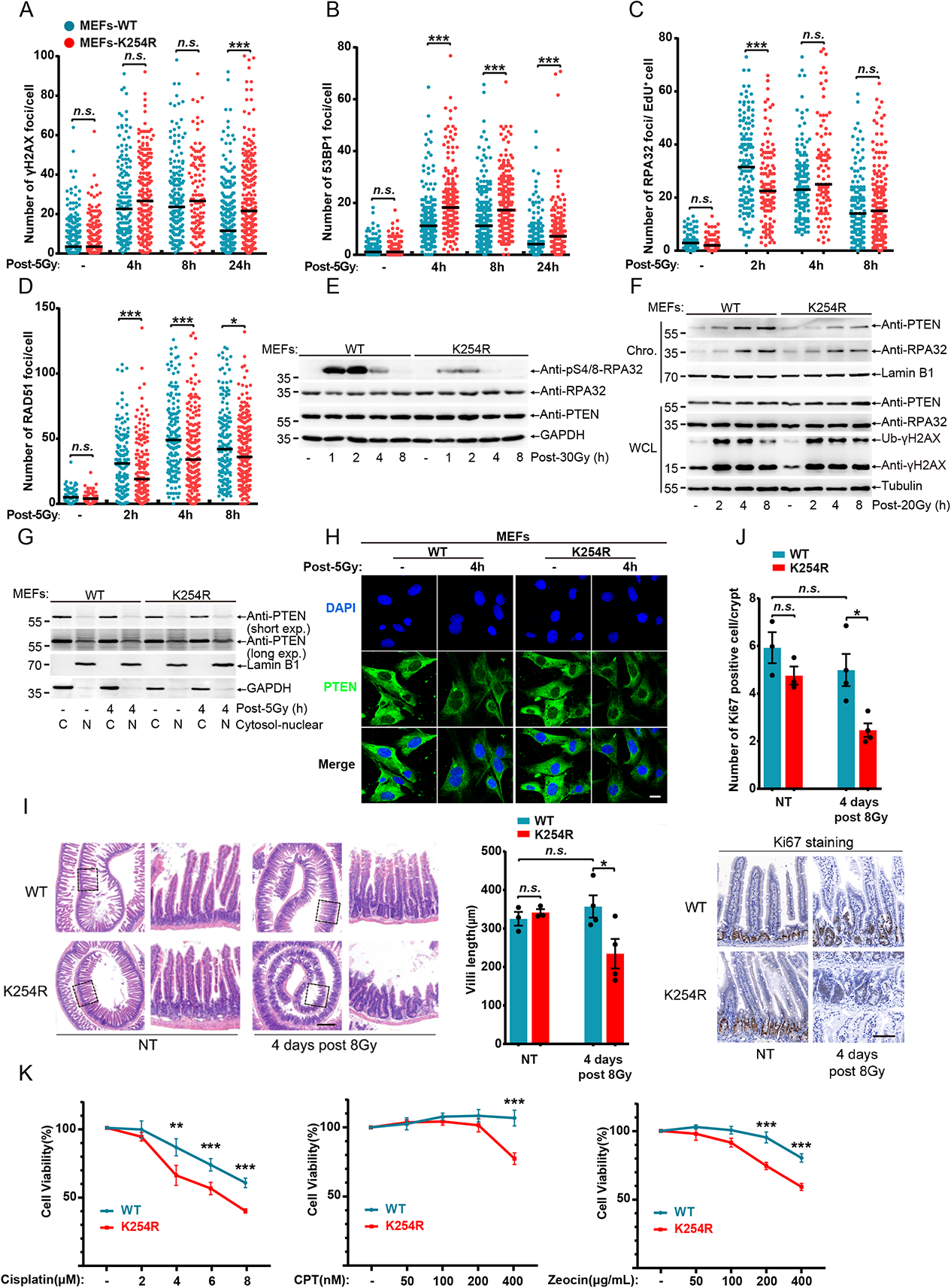
HR repair is impaired by SUMO-deficient PTEN in vivo. (**A-D**) Numbers of 53BP1, γH2AX, RAD51 and RPA32 foci of MEFs post 5 Gy were quantified with FOCO software and presented as dot graph (γH2AX: WT(n(-)=178, n(4h)=185, n(8h)=192, n(24h)=290), K254R(n(-)=270, n(4h)=206, n(8h)=109, n(24h)=282); 53BP1: WT(n(-)=150, n(4h)=183, n(8h)=222, n(24h)=223), K254R(n(-)=229, n(4h)=194, n(8h)=208, n(24h)=149); RPA32: WT(n(-)=111, n(2h)=136, n(4h)=134, n(8h)=135), K254R(n(-)=98, n(2h)=126, n(4h)=97, n(8h)=170); RAD51: WT(n(-)=59, n(2h)=143, n(4h)=152, n(8h)=158), K254R(n(-)=50, n(2h)=153, n(4h)=196, n(8h)=203)). (**E**) Immunoblot of pS4/8-RPA32 of MEFs after treatment with 20 Gy. (**F**) Immunoblot of chromatin loaded PTEN and RPA32 which were separated from MEFs post 20 Gy. (**G**) Nuclear-cytosol separation of MEFs post 5 Gy and detection of PTEN localization with immunoblot. (**H**) Immunofluorescence of PTEN in MEFs after 5 Gy treatment and recovery for 4 h. scale bar, 50 μm. (**I**) Pten^WT^ and Pten^K254R^ mice were treated with 8 Gy whole-body IR. Villus length were quantified at indicated time and representative image of HE stained sections of small intestine were shown (n=3 for NT group, n=4 for IR group). More than 100 villi were assessed from each mouse. scale bar, 300 μm. (**J**) Ki-67 positive cells in intestinal crypts were quantified and representative image of Ki-67 stained sections of small intestine were shown (n=3 for NT group, n=4 for IR group). More than 100 crypts were assessed from each mouse. scale bar, 50 μm. (**K**) Viability of MEFs treated with Cisplatin, Zeocin or CPT were detected with CCK8 and each group were normalized to no treatment group. Unpaired Student’s t-test was used (*p< 0.05, **p< 0.01, ***p< 0.001) and data are shown as mean or mean±s.d. **Figure 6-source data 1.** Western blots of Figure 6.

To validate that K254-SUMOylation is essential for PTEN loading to the chromatin to promote HR repair, we isolated the chromatin for analysis of potential PTEN SUMOylation, showing that a shifted 90-kDa which could be SUMOylated PTEN band, was enhanced in *Pten*^WT^ but not in *Pten*^K254R^ MEFs at 4, 8 h after Zeocin treatment (***Figure 6—figure supplement 1I***). One study reported that K254R mutation may affect the subcellular localization of PTEN (***Bassi et al., 2013***), however our nuclear-cytosol separation results showed there were no any differences between *Pten*^WT^ and *Pten*^K254R^ MEFs under normal condition, even after treatments with IR, CPT or Zeocin (***Figure 6G***, ***Figure 6—figure supplement 1J and K***). Moreover, immunofluorescence staining of PTEN also displayed not much differences in subcellular localization between *Pten*^WT^ and *Pten*^K254R^ under both normal condition and IR treatment (***Figure 6H***).

To compare the protective effect of *Pten*^WT^ and *Pten*^K254R^ mice against DNA damage under physiological condition, villus length and proliferation of intestinal crypts cells, which are highly sensitive to IR due to their rapid turnover rate (***Ma et al., 2019, Xie et al., 2018***), were determined after treatment with whole-body IR. *Pten*^WT^ mice had no significant difference in crypts morphology and villus length between NT (no treatment) and IR (8 Gy) group, which indicated *Pten*^WT^ mice completely recovered, whereas *Pten*^K254R^ mice showed significant destructed crypts and length-shortened villi 4 days after treatment with IR (***Figure 6I***). Similarly, there was no significant difference in numbers of ki67-positive cells between *Pten*^K254R^ and *Pten*^WT^ mice under no treatment, while numbers of ki67-positive cells in intestine crypts of *Pten*^K254R^ mice were less than those of *Pten*^WT^ mice after IR treatment, suggesting reduced proliferating cells of intestine crypts might be due to functional deficiency in DDR (***Figure 6J***). These results proved K254-SUMOylation was essential for PTEN mediating DNA damage repair and irradiation protection *in vivo*. Lastly, we tested the sensitivity of *Pten*^WT^ and *Pten*^K254R^ MEFs to DNA damage reagents, and found that *Pten*^K254R^ MEFs were more sensitive to Cisplatin, CPT and Zeocin than *Pten*^WT^ MEFs in a dose dependent manner (***Figure 6K***). Taken together, above results demonstrate that K254-SUMOylation of PTEN is required for PTEN mediating HR repair in DDR *in vivo*.

### Blocking PTEN SUMOylation pathway sensitizes tumor cells to DNA damage reagents

We firstly determined appropriate concentrations of DNA damage reagents including Cisplatin, Zeocin and CPT for the treatment on DU145-PTEN^WT^ or DU145-PTEN^−/−^ cells, and found that there was little difference in survival colony numbers between DU145-PTEN^−/−^ and DU145-PTEN^WT^ cells when treated with low doses of Cisplatin (0.5 and 1 μM), Zeocin (2 and 5 μg/mL) and CPT (25 and 50 nM), whereas DU145-PTEN^−/−^ cells were much more sensitive to high doses of Cisplatin (1.5 and 2 μM), Zeocin (10 and 15 μg/mL) and CPT (150 nM) than PTEN^WT^ (***Figure 7A-C, Figure 7—figure supplement 1A-C***).

**Figure 7.**
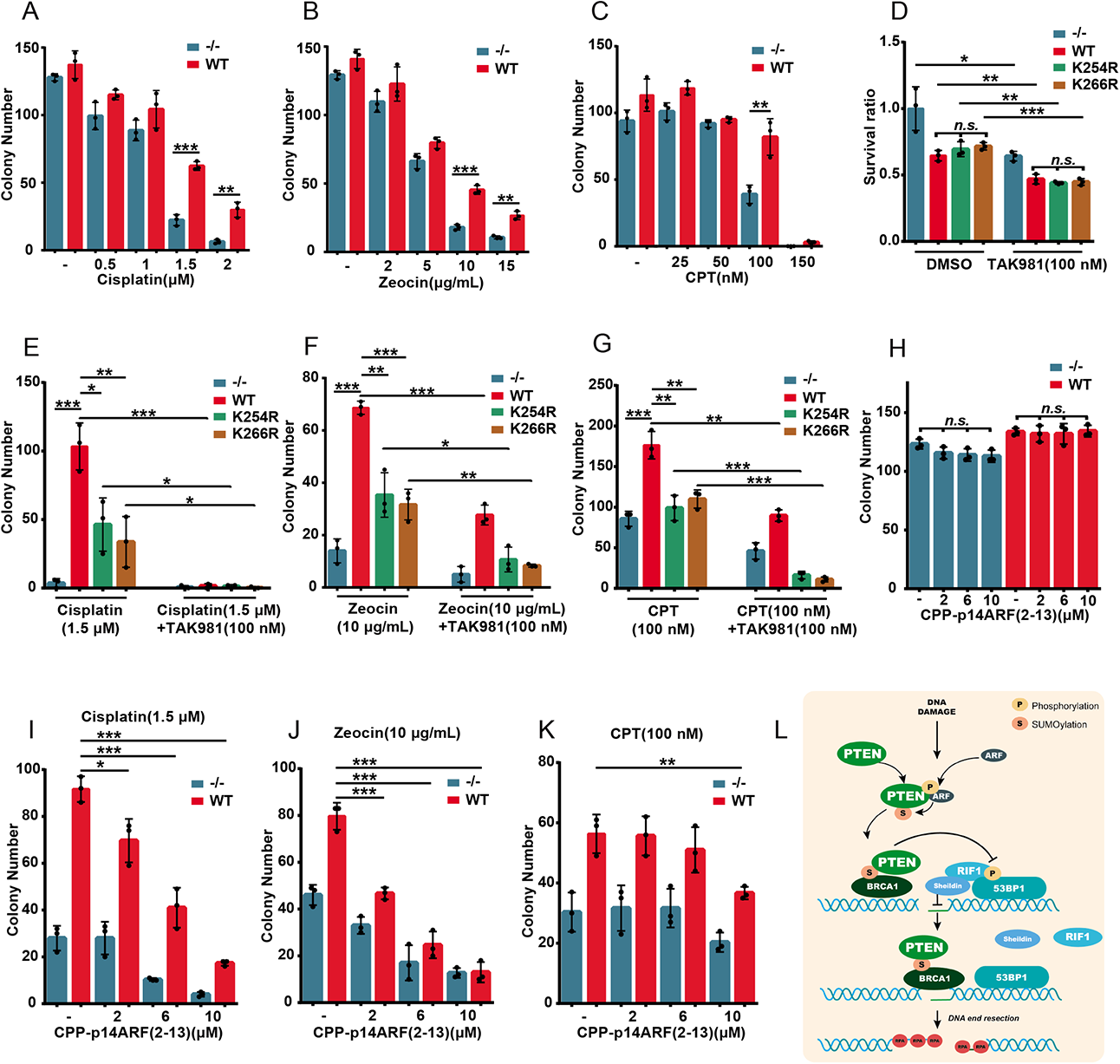
Blocking PTEN SUMOylation pathway sensitizes tumor cells to DNA damage reagents. (**A-C**) DU145-PTEN^−/−^ and PTEN^WT^ cells were treated with or without different doses of Cisplatin for 3 days, Zeocin for 2 days or CPT for 2 days and then cultured for another 7-10 days. Colony number was counted and presented as bar graph (n=3 for each group). (**D**) DU145-PTEN^−/−^, PTEN^WT^, PTEN^K254R^ and PTEN^K266R^ cells were treated with or without TAK981 (100 nM) for 3 days and cultured for another 7-10 days. Total intensity of stained colony was measured by ImageJ and then normalized to DU145-PTEN^−/−^ group without TAK981 treatment (n=3 for each group). (**E-G**) DU145-PTEN^−/−^, PTEN^WT^, PTEN^K254R^ and PTEN^K266R^ cells were treated with Cisplatin (1.5 μM) for 3 days, Zeocin (10 μg/mL) for 2 days or CPT (100 nM) for 2 days or combination with TAK981 (100 nM) and cultured for another 7-10 days. Colony number was counted and presented as bar graph (n=3 for each group). (**H**) DU145-PTEN^−/−^ and PTEN^WT^ cells were treated with different doses of CPP-p14ARF(2-13) for 3 days and cultured for another 7-10 days (n=3 for each group). (**I-K**) DU145-PTEN^−/−^ and PTEN^WT^ cells were treated with combination of Cisplatin (1.5 μM for 3 days), Zeocin (10 μg/mL for 2 days) or CPT (100 nM for 2 days) and different doses of CPP-p14ARF(2-13). Colony number was counted after cultured for another 7-10 days (n=3 for each group). (**L**) A schematic model to show the molecular mechanism of PTEN SUMOylation in HR repair. DNA damage induced phosphorylation of p14ARF enhanced its interaction with PTEN and which subsequently promoted SUMOylation of PTEN. SUMOylated PTEN was recognized and recruited into DNA breaks by BRCA1 and then directly dephosphorylated 53BP1 which helped release of its downstream effectors and DNA end resection. 500 cells were seeded in 12-well plate for all colony assays except (**C**) in which 1000 cells were seeded at the beginning. Unpaired Student’s t-test was used (*p< 0.05, **p< 0.01, ***p< 0.001) and data are shown as mean±s.d from three biological replicates.

Given that SUMOylation signaling is critical for cell survival during DNA damage and a specific SUMO E3 inhibitor is lacking, we tried to assess the cellular sensitivity to DNA damage agents after blocking SUMOylation pathway with SUMO E1 inhibitor, TAK981 (***Langston et al., 2021***), which has been in clinical trial to treat advanced and metastatic solid tumors. In accordance with previous report (***Lightcap et al., 2021***), the total levels of SUMOylation in DU145 cells were inhibited by TAK981 in a dose dependent manner and completely suppressed at the concentration greater than 100 nm (***Figure 7—figure supplement 1D***). We found that TAK981 effectively inhibited the growth/survival of DU145-PTEN^−/−^, -PTEN^WT^, -PTEN^K254R^ and -PTEN^K266R^ cells, although there was no much difference among PTEN^WT^, PTEN^K254R^ and PTEN^K266R^ cells either with or without TAK981 treatment (***Figure 7D, Figure 7—figure supplement 1E***). By assessing combination effects of TAK981 with Cisplatin, Zeocin or CPT, we found that both PTEN^K254R^ and PTEN^K266R^ cells were more sensitive compared with PTEN^WT^ cells (***Figure 7E-G, Figure 7—figure supplement 1E***), which was consistent with our results that PTEN SUMO site mutations of K254R and K266R decreased DNA damage repair efficiency (***Figure 1C and D***). Most strikingly, combination treatment with TAK981 resulted in an additive effect on the sensitivity to Cisplatin, Zeocin or CPT (***Figure 7E-G, Figure 7—figure supplement 1E***). These results indicated that inhibiting the SUMOylation pathway might increase killing efficiency of DNA damage reagents in clinical cancer treatment.

Since p14ARF knockdown inhibited chromatin loading of PTEN and sensitized DU145 cells to DNA damage (***Figure 3K, Figure 3—figure supplement 1H-K***), we further synthesized a small peptide called CPP-p14ARF(2-13), which was a Fitc-labelled cell penetrating peptide (CPP) YGRKKRRQRRR fused by p14ARF(2-13aa)/VRRFLVTLRIRR (Fitc-YGRKKRRQRRRVRRFLVTLRIRR), the main region interacting with and promoting PTEN SUMOylation (***Figure 3F***). We assessed whether CPP-p14ARF(2-13) can interfere PTEN SUMOylation and loading to the chromatin under DNA damage. Indeed, CPP-p14ARF(2-13) dramatically inhibited the interaction between PTEN and p14ARF (***Figure 7—figure supplement 1F***) as well as chromatin loading of PTEN (***Figure 7—figure supplement 1G***) under CPT treatment. Next, we tested whether CPP-p14ARF(2-13) enhances the cell sensitivity to DNA damage. As previously reported that a peptide of p14ARF(1-22aa) inhibited cell proliferation (***Johansson et al., 2008***), so we also detected whether CPP-p14ARF(2-13) influences it. There was no difference in survival colony numbers between PTEN^−/−^ and PTEN^WT^ DU145 cells treated with CPP-p14ARF(2-13) in different doses, suggesting CPP-p14ARF(2-13) did not affect cell proliferation (***Figure 7H, Figure 7—figure supplement 1H***). Interestingly, the cell sensitivity to Cisplatin, Zeocin or CPT treatment was increased by combination with CPP-p14ARF(2-13) (***Figure 7I-K, Figure 7— figure supplement 1H***). The low dose of CPP-p14ARF(2-13) at 2 μM was capable to enhance killing efficiency for Cisplatin and Zeocin treatments to DU145-PTEN^WT^ cells (***Figure 7I and J***), and the high dose at 10 μM also increased killing efficiency for CPT treatment (***Figure 7K***). Furthermore, CPP-p14ARF(2-13) treatment also decreased survival rate of DU145-PTEN^−/−^ cells when combined with these reagents, suggesting CPP-p14ARF(2-13) might block the interaction of p14ARF with other targets besides PTEN. Thus, above results suggest that TAK981 or CPP-p14ARF(2-13) has the potential to enhance the effect of DNA damage reagents in killing tumor cells.

## Discussion

PTMs of PTEN including SUMOylation (***Bassi et al., 2013***), methylation and phosphorylation (***Zhang et al., 2019***) are involved in DDR. Although one study reported that PTEN^K254R^ impaired HR repair efficiency, the underlying molecular mechanism remained largely elusive (***Bassi et al., 2013***). Our data suggested that DNA damage induced PTEN SUMOylation in a time dependent manner other than rapid activation like phosphorylation. Both K254 and K266 of PTEN could be conjugated with SUMO1, but one single mutation was enough to reduce DNA damage-induced PTEN SUMOylation. Interestingly, the mutation K254R, but not K266R, suppressed DNA damage-triggered PTEN chromatin loading. It is an open question how K266-SUMOylation of PTEN to participate in DDR. Given that K266 has also been identified as a ubiquitination site (***He et al., 2021***), it might exist a crosstalk between SUMOylation and ubiquitination of PTEN during DDR. Moreover, SUMOylation of PTEN on K266 promotes its association with cell membrane and which in turn inhibits PI3K-AKT signaling (***Huang et al., 2012***). As PI3K-AKT pathway can be also activated by DNA damage stimuli and regulated function of DDR related proteins(***Golding et al., 2009, Hu et al., 2017, Jia et al., 2013, Plo et al., 2008***), K266-SUMOylation of PTEN might also promote DDR through the PI3K-AKT pathway. Additionally, the mutation C124S remarkably enhanced PTEN nuclear localization and chromatin loading. Due to PTEN interacting with RAD51 and RPA32 (***He et al., 2015, Wang et al., 2015***), this might explain why PTEN^C124S^ decreased HR repair efficiency but not affected the chromatin loading of RAD51 and RPA32. It remains to be further explored how C124S influences PTEN localization and whether the chromatin trapped PTEN^C124S^ has other additive side effects on DNA damage repair.

Several SUMO E3 ligases including PIAS1, PIAS4, CBX4 and ZNF451 play an important role in regulation of SUMOylation signal transduction in response to DNA damage (***Galanty et al., 2009, Soria-Bretones et al., 2017, Tian et al., 2021***). Here, we identified an atypical SUMO E3 ligase, p14ARF, responsible for PTEN SUMOylation under DNA damage such as IR, CPT and Zeocin through their interaction, which was enhanced by DNA damage-induced phosphorylation of p14ARF. UV stress disrupts the p14ARF-B23 interaction in the nucleolar, resulting in a transient translocation of p14ARF to the nucleoplasm (***Lee et al., 2005***). In addition, ATM, a main DDR kinase, promotes the release of p14ARF from the nucleus and subsequent degradation after doxorubicin treatment (***Velimezi et al., 2013***). So, the subnuclear distribution and protein-protein interactions of p14ARF are critical for its function in promoting PTEN SUMOylation after DNA damage.

As for the role of PTEN protein phosphatase in HR repair, the conclusions from two different groups are contradictory (***Ma et al., 2019, Zhang et al., 2019***). Our results supported that PTEN protein phosphatase was necessary for HR repair. In addition to γH2AX as a substrate of PTEN during DDR (***Zhang et al., 2019***), we identified a new target, 53BP1, which was directly dephosphorylated by PTEN in cells and *in vitro*. More interestingly, PTEN selectively dephosphorylated pT543-53BP1 other than pS25-53BP1 in cells. It has been early reported that phosphorylation at 7 S/TQ sites in the N-terminal region of 53BP1 is responsible for its recruitment of RIF1, whereas phosphorylation at the other 8 S/TQ sites of 53BP1 is essential for PTIP accumulation at DNA-break sites (***Callen et al., 2013***). And interestingly, as long as one site of 7 S/TQ is phosphorylated, it is sufficient for 53BP1 recruiting RIF1 (***Isono et al., 2017***). However, most recently one study reported that RIF1 is recruited to IR-induced foci by recognizing three related phosphorylated epitopes on 53BP1 (***Setiaputra et al., 2022***). Thus, PTEN might also target phosphorylation at other sites responsible for RIF1 recruitment, besides pT543-53BP1 that is one of 7 S/TQ, during DDR.

As known that BRCA1 is critical for HR repair by against 53BP1 posted barrier through serval molecular mechanisms, including inhibition of 53BP1 phosphorylation (***Isono et al., 2017***). Our results supported that PTEN was a downstream effector of BRCA1 and PTEN SUMOylation was required for their interaction, which was efficiently decreased by double mutation of K254/266R. One SIM located in the N-terminal of BRCA1 was essential for recognition of SUMO1 conjugated to PTEN, by which PTEN was subsequently recruited by BRCA1 to DNA-break sites. This might partially explain why that the deletion of *exon11* of BRCA1 resulted in loss of BRCA1 function in inhibition of phosphorylation of 53BP1 (***Nacson et al., 2020, Nacson et al., 2018***).Moreover, as 53BP1 is pro-choice for DNA breaks and BRCA1 complex can relocate into the core of 53BP1 foci at S/G2 in a time dependent manner (***Chapman et al., 2012, Zimmermann et al., 2014***), so we speculate that after entering the core, BRCA1 recruits PTEN to directly dephosphorylate 53BP1, thus releasing downstream effectors of 53BP1 such as RIF1 and PTIP, which further facilitates DNA end resection and ongoing of HR repair.

The *in vivo* results from SUMO-deficient *Pten*^K254R^ mice validated that PTEN SUMOylation promoted HR repair. Significantly, we did not observe the mutation K254R influence PTEN nuclear localization (***Bassi et al., 2013***), by *in vivo* and *in vitro* results, which is consistent with our early report (***Huang et al., 2012***). However, our data strongly demonstrated that SUMOylation of PTEN was induced by several DNA damage agents and this sub-pool of SUMOylated PTEN was tightly associated with chromatin *via* BRCA1.

Since we proved that activation of SUMOylation pathway was essential for correct DNA damage repair, so the combination of SUMOylation inhibitor with DNA damage reagents might enhance sensitivity of tumor cells to chemotherapy. As expectedly, the SUMOylation inhibitor TAK981 remarkedly increased killing efficiency of DNA damage reagents. Furthermore, the small peptide CPP-p14ARF(2-13) also suppressed DNA damage-induced chromatin loading of PTEN and sensitized tumor cells to chemotherapy.

In summary, our study uncovers a new mechanism that SUMOylated PTEN promotes HR repair through dephosphorylation of 53BP1 (***Figure 7L***). In response to DNA damage, p14ARF as a SUMO E3 ligase is phosphorylated to enhance the interaction with PTEN in the nucleus, which subsequently promotes PTEN SUMOylation. Then SUMOylated PTEN is recognized and recruited to the chromatin near DSB by the N-terminal SIM of BRCA1. This pool of PTEN relieves HR repair barrier posted by 53BP1 through directly dephosphorylating 53BP1, promoting HR repair. Blocking PTEN SUMOylation pathway by TAK981 and CPP-p14ARF(2-13) sensitizes tumor cells to DNA damage reagents. Thus, our study elucidated a new molecular mechanism of the key role of PTEN in HR repair during DDR, which may provide a new strategy for clinical cancer therapy.

## Materials and methods

### Cell culture, transfection and lentiviral infection

HEK293T, HEK293FT, DU145, HeLa, H1299 and MEFs were cultured in DMEM supplemented with 10% FBS and 1% 100 U of penicillin, and 100 μg/mL streptomycin (Yeasen). PC3 was cultured in RPMI1640 supplemented with 10% FBS and 1% 100 U of penicillin, and 100 μg/mL streptomycin (Yeasen). DU145-PTEN^−/−^ was generated with CRSPR/Cas9. U2OS-DR-GFP was a gift from Dr. Daming Gao (***Xie et al., 2018***). *Pten^WT^* and *Pten^K254R^* MEFs were obtained from wild-type and *Pten^K254R^* knock-in mice, respectively, and then immortalized with SV40-LT at passage three. Plasmids and siRNA transfection were carried out with PEI for HEK293T, HEK293T*^Senp−/−^* and HEK293FT and Lipo2000 (Invitrogen) for other cells following the manufacturer’s protocol. Packaging lentiviral and subsequent infection of all cell lines were carried out according to protocol in our laboratory.

### Antibodies, reagents, plasmids, siRNA, shRNA and sgRNA

Antibodies used in this study were listed in Supplementary file 1A. PTEN cDNA was subcloned into pCMV-Flag, pEF-5HA, pEGFP-C1, pCD510B and pGEX-4T-1 vectors. shRNAs of BRCA1, PTEN and p14ARF were designed and cloned into the vector pLKO.1. RFP-PCNA was a gift from Prof. Pumin Zhang (***Ha et al., 2017***). Flag-53BP1 and Myc-BRCA1 were gifts form Prof. Xingzhi Xu (***Peng et al., 2015***). pCBASceI, EJ5-GFP and DR-GFP were purchased from Addgene. p14ARF was cloned into pGEX-4T-1, pEYFP-N1 and pCD513B vectors. PCR-mediated site-directed mutagenesis and truncated proteins were performed according to standard procedures to create the PTEN, p14ARF, 53BP1 and BRCA1 mutants. All clones were sequenced to confirm the desired mutations. siRNAs targeting BRCA1 were synthesized by GenePharma. Two sgRNAs were designed to knockout PTEN. In brief, sgRNA was insert into LentiCRISPR v2 and delivered into DU145 cells, each single clone was selected and cultured. PTEN knockout clones were identified and used in our study. A list of the sequence information for the shRNAs, siRNAs and sgRNAs was provided in Supplementary file 1B.

### Immunoblot and denatured immunoprecipitation for SUMOylation detection

Cells were washed once with PBS and lysed in lysis buffer (50 mM Tris-HCl pH 7.4, 150 mM NaCl, 1 mM DTT, 1 mM EDTA, 1% NP-40, complete protease inhibitor cocktail (Roche) and 20 mM N-ethylmaleimide) on ice for 30 min, then lysates were sonicated and centrifuged at 12,000 g for 30 min. Protein concentrations were quantified with BCA kit (Thermo-Fisher). Equivalent amounts of protein (1-2 mg) were incubated with 1 μg indicated antibody and 20 μL protein A/G beads (#IP05, Calbiochem) at 4 ℃ overnight. Beads were collected with centrifugation and washed with lysis buffer for 5 times, and then boiled with 2x protein loading buffer before analysis by SDS-PAGE.

Denatured immunoprecipitation for SUMOylation detection was carried out as previously described with several modifications (***Yu et al., 2009***). Briefly, cells were lysed with SUMO lysis buffer (62.5 mM Tris pH 6.8, 2% SDS), sonicated, and boiled after addition of 1 mM DTT. The lysis was centrifuged at 12,000 g for 15 min. Supernatant was transferred into new EP tube and diluted to a final concentration of 0.1% SDS with lysis buffer. Equivalent amounts of protein were incubated with indicated antibodies (anti-PTEN, anti-HA and anti-Flag) and protein A/G beads overnight at 4 ℃. Beads were washed with lysis buffer containing 300 mM NaCl and 0.1% SDS for 5 times and boiled with 2x protein loading buffer before analysis by SDS-PAGE.

### Ni^2+^-NTA pull down for SUMOylation assay

For detection of PTEN SUMOylation during DNA damage repair, 293T cells were transfected with His-SUMO1 and indicated plasmids for 48 h. CPT (#S1288, Selleck) and Zeocin (#60216ES80, Yeasen) were used to induce DNA damage for 1 h. cells were harvested at indicated time, 10% cells were used as input. SUMO-PTEN were pulled down with Ni^2+^-NTA beads (#30210, Qiagen) and analyzed with SDS-PAGE as previous described (***Huang et al., 2012***).

### Immunofluorescence

Cells were seeded on glass coverslips. After treatment with various DNA damage stimuli, cells were washed with PBS and then fixed with 4% (w/v) paraformaldehyde in PBS for 15 min at room temperature. When staining RPA32, cells were pre-extracted with cold 0.5% Triton X-100 in PBS for 3 min before fixing. After fixing, cells were permeabilized with 0.5% (v/v) Triton X-100 in PBS for 60 min and blocked with 5% BSA in PBS for 60 min at room temperature. Generally, cells were then incubated with the primary antibody diluted in PBS-BSA overnight at 4 ℃. Cells were washed 3 times with PBST and then incubated with secondary antibodies diluted in PBS-BSA supplemented with 2 µg/ml of Hoechst 33342 (#62249, Thermo-Fisher) to stain DNA for 1 h at room temperature. Cells were washed 3 times with PBST and then the coverslips were mounted onto glass slides with Prolong Gold mounting agent (#P36931 Thermo-Fisher). Confocal images were taken with a LSM710 or Zeiss LSM880 laser-scanning confocal microscope.

For the discrimination of cells at S stage, cells were pre-incubated with 10 µM EdU (#C0081S, Beyotime) for 30 min before DNA damage induction. Click-It reaction were carried out as manufacture’s protocol to staining EdU positive cells. After this, cells were used for further immunostaining. Intensity and number of DNA damage induced foci were counted with FOCO software (***Lapytsko et al., 2015***).

### HR repair assay

To determine the efficiency of HR-mediated DSB repair in cells expressing different form of PTEN with mutation, we firstly generated stably PTEN knockdown U2OS-DR-GFP cells with shRNAs. Then PTEN-WT, C124S, G129E, K254R, K266R were re-expressed in those cells. To increase RFP-I-Sce1 expression efficiency, RFP-I-Sce1 was subcloned into the vector CD510B. Pseudo-lentivirus expressing RFP-I-Sce1 was used to infect U2OS-DR-GFP cells. After 48-72 h, cells were collected for FACS analysis. Percentage of RFP and GFP positive cells were quantified. HR efficiency was quantified as (GFP^+^/RFP^+^) *100%.

### Laser micro-irradiation

Generation of localized DNA damage by laser was done as previously described (***Ha et al., 2017***). Briefly, cells were seeded in a live-cell imaging culture dish, transfected with GFP-PTEN and RFP-PCNA and cultured for 48 h. 2 μg/mL Hoechst 33342 was used to pre-sensitize cells for 10 min before laser micro-irradiation. For micro-irradiation, the cell dish was mounted on the stage of a Leica SP8 microscope at 37 ℃. 405 nm UVA focused through a 63x 1.4NA oil objective was used to induce localized DNA damage. Laser power was set to 50% and iterations were set to 50 times. Time-lapse imaging of recruitment of GFP- or RFP-tagged proteins to DNA damage site was captured every 30 sec after micro-irradiation with 488 and 561 nm laser.

### GST pull-down

GST-fused protein was expressed in *E. coli* BL21 and affinity-purified with GST beads (#17-0756-01, GE Healthcare Life Sciences). 293T cells overexpressing indicated proteins were treated with DNA damage reagents and lysed. And then cell lysates were incubated with above GST-fused protein beads overnight. Beads were collected and washed 5 times, followed by Western blotting analysis.

### *In vitro* dephosphorylation assay

For measuring dephosphorylation of 53BP1 by PTEN, the GST-fused PTEN^WT^, PTEN^C124S^, PTEN^G129E^, and PTEN^G129R^, as well as GST protein were purified from *E. coli* BL21. pT543-53BP1 peptide was synthesized by Top-peptide Biotech Co., Ltd (Shanghai, CN). Full-length Flag-53BP1 and GFP-53BP1 were purified from 293T cells after Zeocin treatment. The dephosphorylation assays were performed in phosphatase assay buffer (20 mmol/L HEPES, pH 7.2, 100 mmol/L NaCl, and 3 mmol/L DTT). The reactions were incubated at 37°C for 60 min with or without the addition of recombinant GST-fused PTEN^WT^, PTEN^C124S^, PTEN^G129E^, or PTEN^G129R^, as well as GST protein as negative control, and then were stopped by adding 2× SDS loading buffer for immunoblotting analysis.

### Cellular fractionation

MEFs, DU145, H1299 cells cultured with 90%-100% confluence were harvested after treatment with DNA damage reagents and recovery for indicated time. Extraction of cytoplasmic and nuclear proteins was performed using the Nuclear/Cytosol Fractionation Kit (BioVision) according to its instruction.

Separation of chromatin associated proteins was performed as previously described with minor modification (***Mendez et al., 2000***). Briefly, cell pellets were washed with cold PBS and then incubated with buffer A (10 mM pH 7.9 HEPES, 10 mM KCl, 1.5 mM MgCl2, 0.34 M sucrose, 10% glycerol, 1 mM DTT, protease inhibitors, 0.1% Triton-X100) for 10 min on ice. Cell pellets were collected with centrifugation and washed twice with buffer A. Next, cell pellets were gently resuspended in buffer B (3 mM EDTA, 0.2 mM EGTA, 1 mM DTT, protease inhibitors) and incubated for 30 min on ice. Then cell pellets were collected by centrifugation and lysed in 2% SDS as chromatin associated proteins. When detecting proteins tightly associated with chromatin, we pipetted cell pellets with buffer B harshly until sticky chromatin pellets were visible after incubation with buffer A. After another 30 min incubation in buffer B, chromatin pellets containing tightly associated proteins was collected by centrifugation and lysed with 2% SDS.

### Cell viability and colony formation assay

For cell viability assay, cells were counted and 10000 cells were seeded into 96 well plate. After 24 h, DNA damage reagents were added and cultured for another 3 days. CCK8-kit was used to detect cell viability and all quantitative results were normalized to non-treatment group. For colony formation assay, 500 or 1000 cells were seeded into a 12-well plate. After 24 h, CPT and Zeocin were added for 48 h and replaced with fresh medium. Cisplatin was added into medium for 72 h and replaced with fresh medium. All culture medium was changed every 3 days until colony was visible. Colonies were washed, fixed and stained with 0.1% crystal violet overnight. Visible colonies were counted and analyzed between groups with ImageJ.

### Mouse model and IHC

All animal studies were conducted with the approval and guidance of Shanghai Jiao Tong University Medical Animal Ethics Committees. *Pten*-K254R and *Pten*-K266R knockin mice were generated by BRL medicine company with CRISPR-Cas9 and homozygous mice were verified with PCR sequencing. For IHC, 4-month-old male mice were chosen and subjected to whole body irradiation (IR) with 8 Gy. Mice were sacrificed at day 4 post IR, small intestines were used for histological analysis. HE staining were used to quantify villi length, and Ki67 staining was used to identify proliferating cells in small intestines.

### Statistical analysis

Group data are presented as mean with or without ± s.d. The statistical significance between experimental groups was determined by Student’s t-test (two tailed and unpaired). p<0.05 was considered to be significant (*n.s.*, not significant; *0.01<P<0.05, **0.001<P<0.01 and ***P<0.001). If not specified, analysis was performed with GraphPad Prism 8.

## Acknowledgments.

This work was supported by grants from China’s National Key R&D Programmes (NKP) (No. 2019YFE0110600), the National Natural Science Foundation of China (82230100, 81630075, 32271310, 82103082, 81721004) and Shanghai Science and Technology Commission (20JC1410100).

## Author contributions

J.Y., J.He. and R.D. conceived and designed the study. J.H., Y.G., R.D. and L.L. performed most of the experiments. C.H., R.C., Y.W., X.Z. and J. Huang helped with experiments and provided technical support. J.Z., G.C, and J.C. offered constructive suggestions. J.Y., J.He. Y.G. and R.D. analyzed the data. J.Y., J.He. and X.Z. wrote the manuscript. All authors read and approved the final manuscript.

## Data availability

All data generated or analysed during this study are included in the manuscript and supporting files.

## Figure legends

**Figure 1-figure supplement 1.**
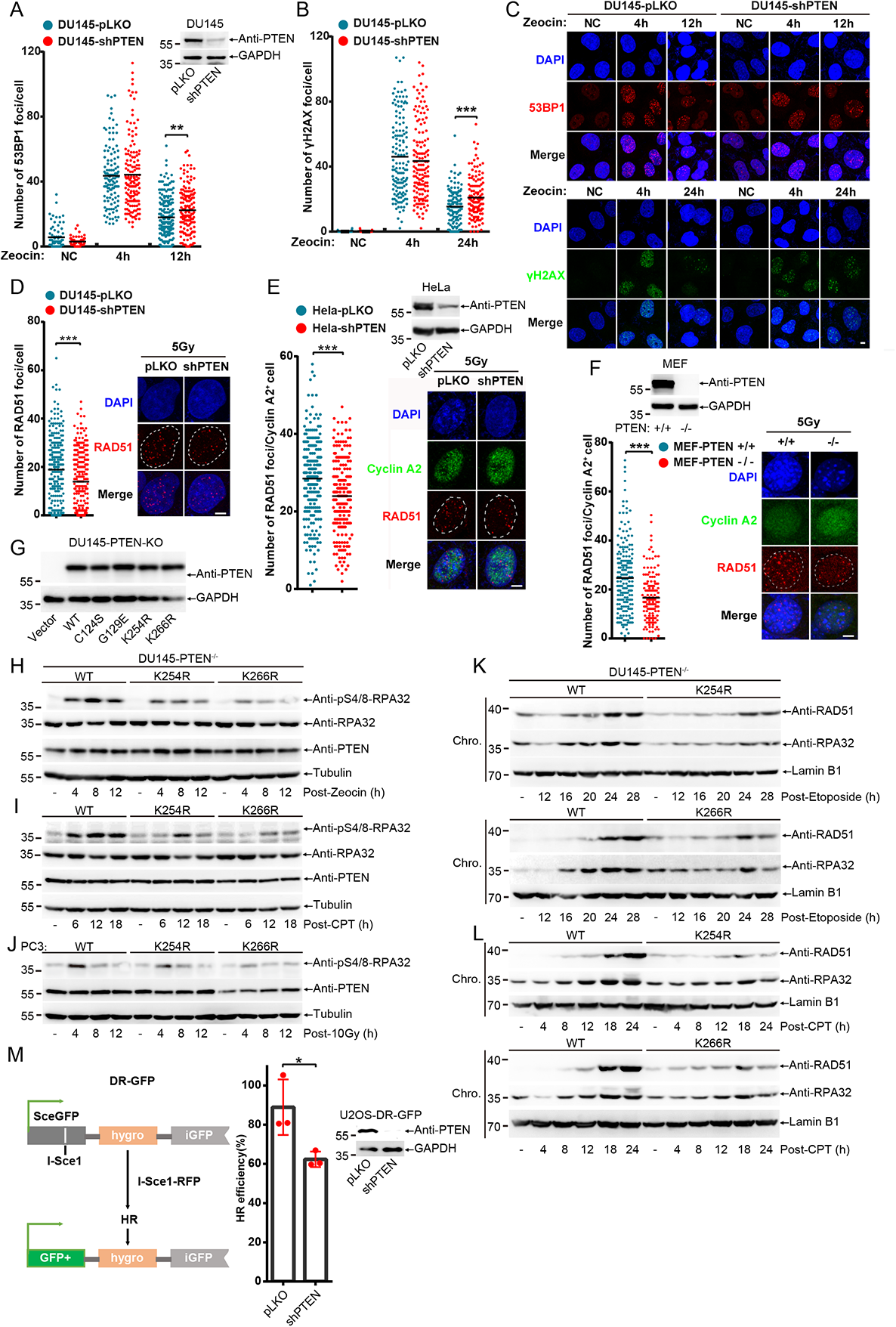
PTEN promotes HR repair through enhancing DNA end resection. (A-C) Quantification of 53BP1 and γH2AX foci in DU145 cells after treatment with Zeocin (200 μg/mL) for 1 h and recovery for indicated time (53BP1: DU145-pLKO(n(NC)=74, n(4h)=121, n(12h)=205), DU145-shPTEN(n(NC)=66, n(4h)=143, n(12h)=154); γH2AX: DU145-pLKO(n(NC)=36, n(4h)=142, n(24h)=134), DU145-shPTEN(n(NC)=63, n(4h)=141, n(24h)=130)). Inset: immunoblot of PTEN knockdown efficiency in DU145 cells. Representative immunofluorescence images were shown. scale bar, 20 μm. (D-F) Quantification of RAD51 foci in DU145, HeLa and MEF cells after treatment with 5 Gy and recovery for 6 h (DU145-pLKO(n=259), DU145-shPTEN(n=266); HeLa-pLKO(n=235), HeLa-shPTEN(n=203); MEF-PTEN^+/+^(n=196), MEF-PTEN^−/−^(n=144)). Representative immunofluorescence images were shown. Inset: immunoblot of PTEN knockdown or knockout efficiency in HeLa and MEF cells. scale bar, 20 μm. (G) Immunoblot of PTEN in DU145-PTEN^−/−^ cells stably re-expressing PTEN^WT, C124S, G129E, K254R and K266R^. (H-I) Immunoblot of pS4/8-RPA32 level in DU145-PTEN^−/−^ cells stably re-expressing PTEN^WT, K254R and K266R^ after treatment with CPT (20 μM) or Zeocin (200 μg/mL) for 1h and recovery for indicated time. (J) Immunoblot of pS4/8-RPA32 level in PC3 cells stably re-expressing PTEN^WT, K254R and K266R^ after treatment with 10 Gy and recovery for indicated time. (K,L) Immunoblot of chromatin associated RAD51 and RPA32 proteins separated from DU145 cells after treatment with Etoposide (30 μM) or CPT (20 μM) for 1 h and recovery for indicated time. (M) Left panel describes how DR-GFP reporter works. HR efficiency were quantified in U2OS-DR-GFP and U2OS-DR-GFP-shPTEN cells after transfection of I-Sce1 for 48-72h at right panel (n=3 for each group). Inset: immunoblot of PTEN knockdown efficiency in U2OS-DR-GFP cells. Unpaired Student’s t-test was used (*p< 0.05,**p< 0.01, ***p< 0.001) and data were shown as mean or mean±s.d. **Figure 1-figure supplement 1-source data 1. Western blots of Figure 1-figure supplement 1.**

**Figure 2-figure supplement 1.**
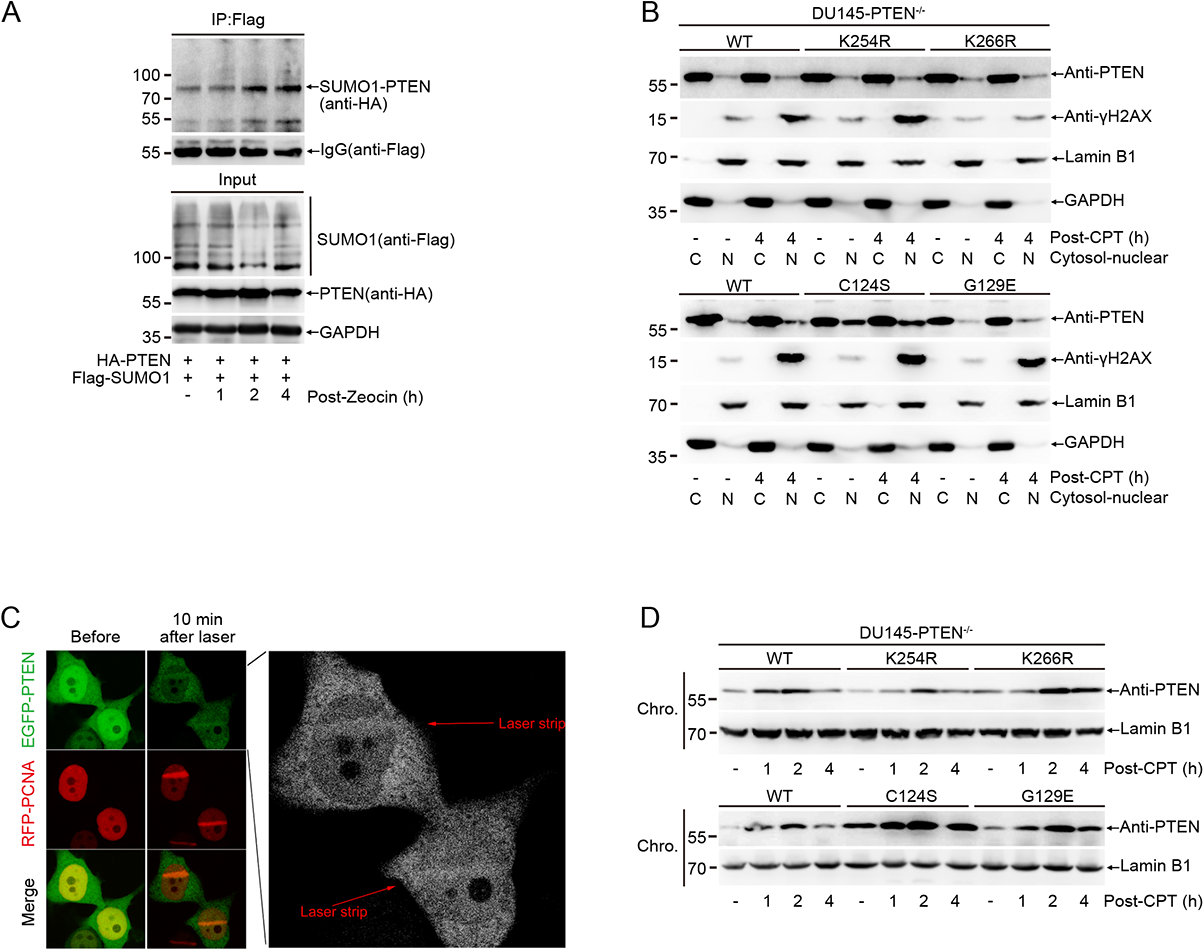
DNA damage promotes PTEN chromatin loading by inducing its SUMOylation. (A) Denatured Co-IP were performed to analyze SUMOylation of PTEN in 293T cells transfected with Flag-SUMO1 and HA-PTEN after treatment with Zeocin (400 μg/mL). (B) PTEN localization was detected with immunoblot after Nuclear-Cytosol separation in DU145-PTEN^−/−^ cells stably re-expressing PTEN mutants after treatment with CPT (20 μM) for 1 h and recovery for 4 h. (C) 293T^senp−/−^ cells transfected with PCNA-RFP and EGFP-PTEN were treated with laser micro-irradiation. Images were taken before and 10 min post laser micro-irradiation. PCNA-RFP were used as indicator of DNA damage location. (D) Chromatin loading of PTEN was detedcted with immunoblot in DU145-PTEN^−/−^ cells stably re-expressing PTEN mutants after treatment with CPT (20 μM) for 1 h and recovery for indicated time. **Figure 2-figure supplement 1-source data 1. Western blots of Figure 2-figure supplement 1.**

**Figure 3-figure supplement 1.**
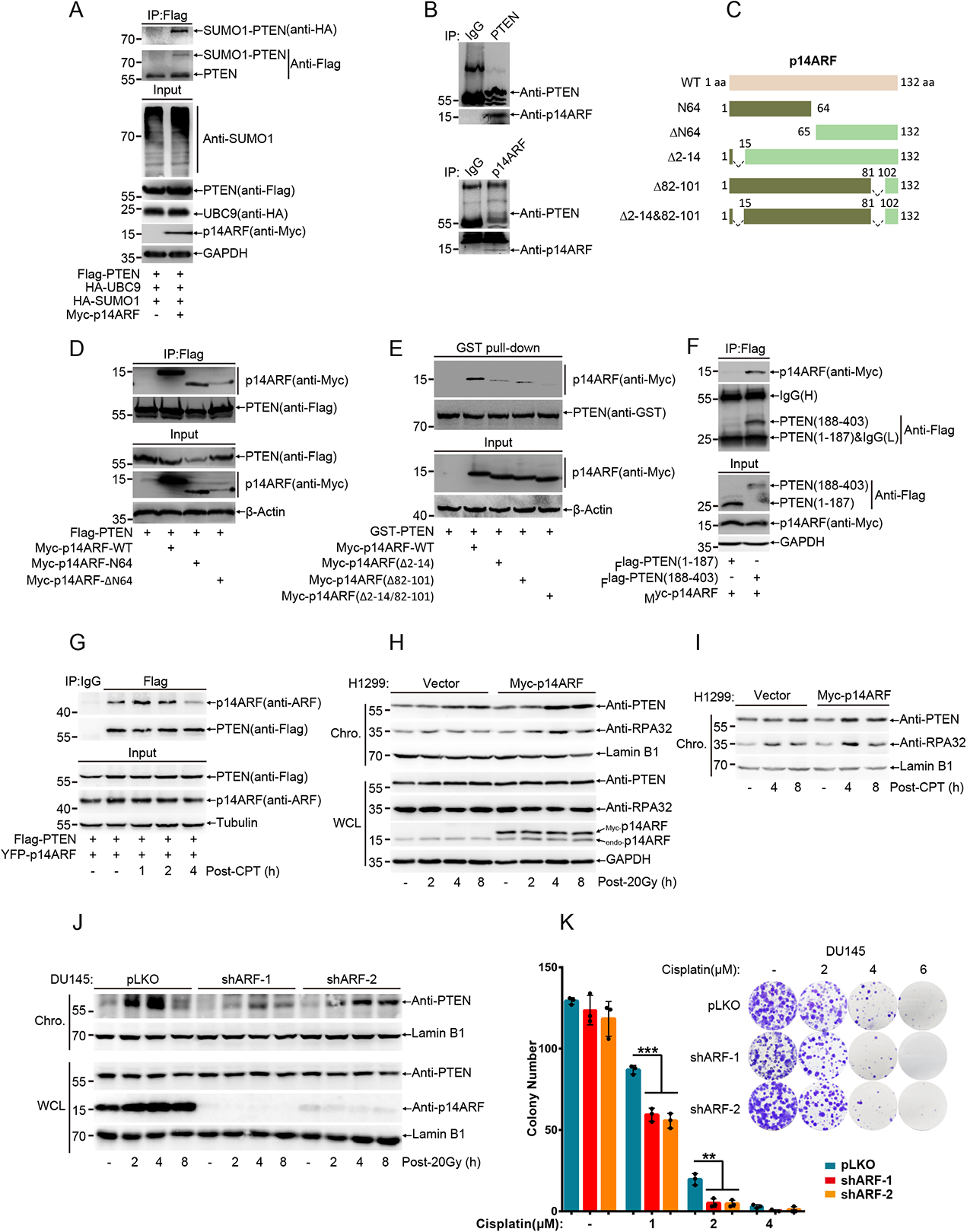
p14ARF is a novel SUMO E3 ligase to mediate PTEN SUMOylation during DDR. (A) SUMOylated PTEN was detected with IP in 293T cells overexpressed indicated plasmids. (B) Reciprocal endogenous interaction between PTEN and p14ARF were detected with Co-IP in 293T cells. (C) Schematic structure of truncated p14ARF. (D) Truncated Myc-p14ARF and Flag-PTEN were transfected into 293T cells for 48 h, interaction domain between PTEN and p14ARF were identified with Co-IP. (E) GST-PTEN was purified from BL21 and incubated with lysis of 293T cells transfected with different truncated Myc-p14ARF, PTEN bound truncated Myc-p14ARF was identified with GST pull-down and immunoblot. (F) Truncated Flag-PTEN and Myc-p14ARF were transfected into 293T cells for 48 h, domain of PTEN responsible for interacting with p14ARF were identified with Co-IP. (G) Interaction between PTEN and p14ARF after treatment with CPT (20 μM) for 1 h and recovery for indicated time was identified with Co-IP. (H, I) Chromatin loaded PTEN were separated from H1299 stably expressing Vector or Myc-p14ARF after treatment with 20 Gy or CPT (20 μM) for 1 h and recovery for indicated time. (J) Immunoblot of chromatin loaded PTEN which separated from DU145-pLKO, shARF-1 and shARF-2 cells after treatment with 20 Gy and recovery for indicated time. (K) Colony survival of DU145-pLKO, shARF-1 and shARF-2 cells after treatment with different doses of Cisplatin. Colony number was counted and shown as mean±s.d. from three independent experiments at left panel. Representative colony images were shown at right panel. Unpaired Student’s t-test was used (**p< 0.01, ***p< 0.001). **Figure 3-figure supplement 1-source data 1. Western blots of Figure 3-figure supplement 1.**

**Figure 4-figure supplement 1.**
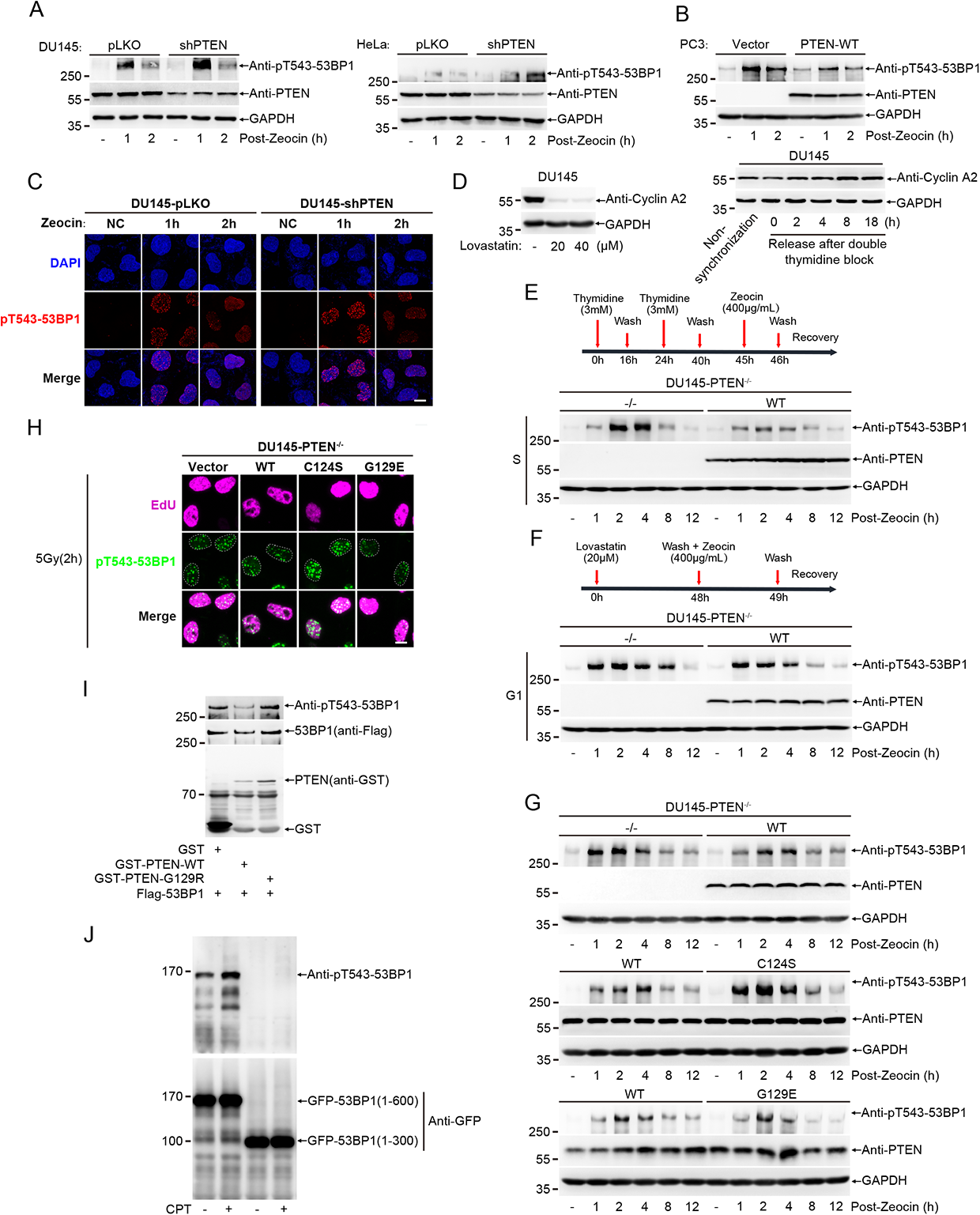
PTEN relieves HR barrier posted by 53BP1 through directly dephosphorylating pT543-53BP1. (A) Immunoblot of pT543-53BP1 level in DU145 and HeLa cells stably expressing pLKO or shPTEN after treatment with Zoecin (200 μg/mL) for 1 h and recovery for indicated time. (B) Immunoblot of pT543-53BP1 level in PC3 cells stably expressing Vector or PTEN^WT^ after treatment with Zoecin (200 μg/mL) for 1 h and recovery for indicated time. (C) Representative images corresponding to Fig. 4B. (D) Cell cycle were synchronized to G1 with lovastatin or to S phase with double thymidine block in DU145 cells. Immunoblot of Cyclin A2 was shown to detect the synchronization efficiency. (E) Schematic illustration of S phase synchronization and Zeocin addition was shown at upper panel. Immunoblot of pT543-53BP1 level in DU145 cells after S synchronization was shown at lower panel. (F) Schematic illustration of G1 phase synchronization and Zeocin addition was shown at upper panel. Immunoblot of pT543-53BP1 level in DU145 cells after G1 synchronization was shown at lower panel. (G) Immunoblot of pT543-53BP1 level in DU145 cells after Zeocin (400 μg/mL) treatment for 1 h and recovery for indicated time. (H) Representative images corresponding to Fig. 4H. (I) Immunoblot of total pT543-53BP1 level after *in vitro* dephosphorylation assay. Full-length Flag-53BP1 were immunoprecipitated from 293T cells after treatment with CPT (20 μM) and recovery for 1 h and then reacted with purified GST-PTEN^WT^ and PTEN^G129R^ (a dual phosphatase deficient mutant) which were purified from BL21. (J) Immunoblot of pT543-53BP1 from 293T cells which were transfected GFP-53BP1(1-300) or GFP-53BP1(1-600) after treatment with CPT (20 μM) for 1 h and recovery for 1 h. **Figure 4-figure supplement 1-source data 1. Western blots of Figure 4-figure supplement 1.**

**Figure 5-figure supplement 1.**
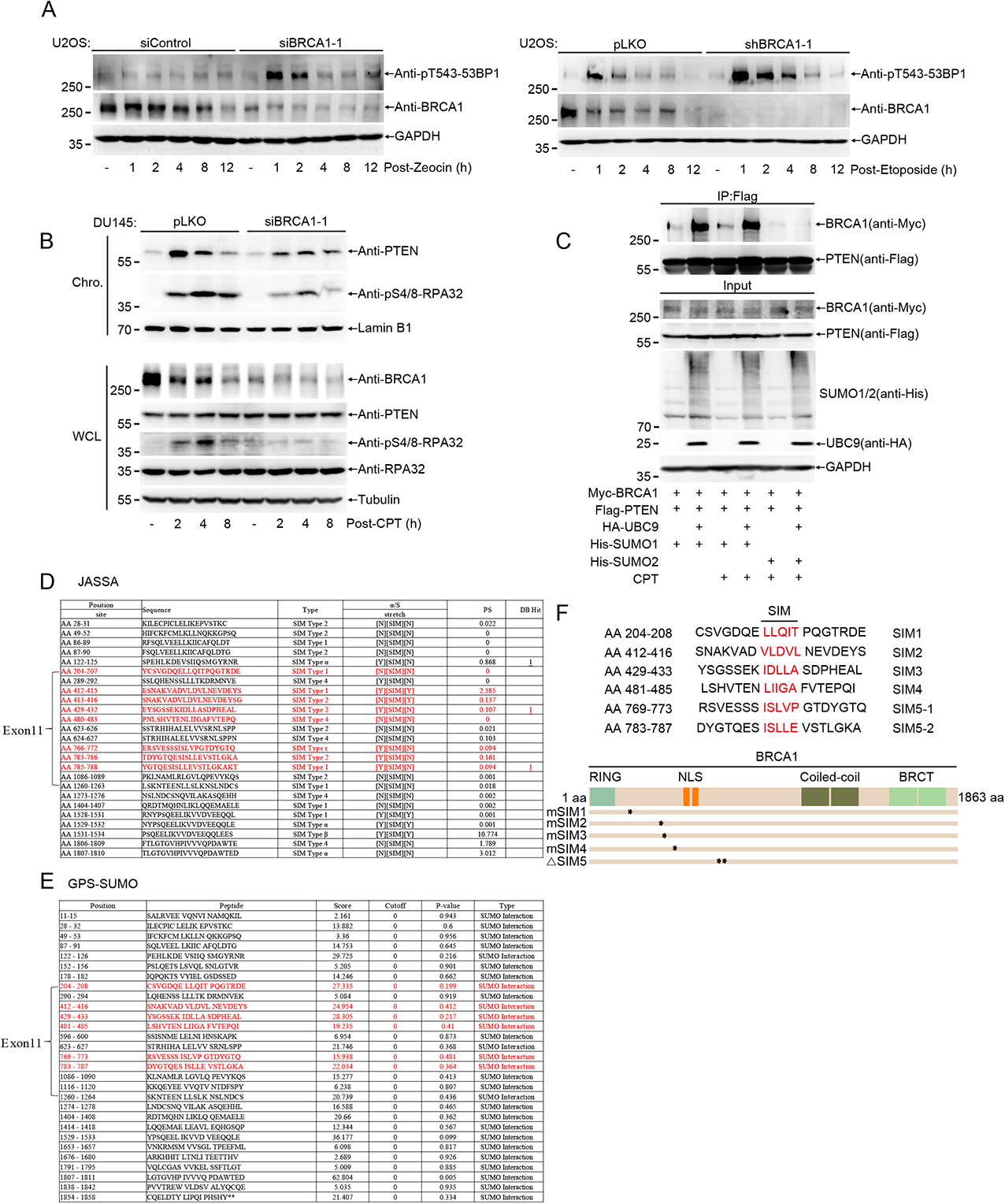
PTEN chromatin loading is mediated by BRCA1 recruiting SUMOylated PTEN via its N-terminal SIM. (A) pT543-53BP1 were detected with immunoblot in U2OS cells in which BRCA1 was knocked down with siRNA or shRNA after treatment with Zeocin (400 μg/mL) or Etoposide (30 μM) for 1 h and recovery for indicated time. (B) Chromatin loaded PTEN was detected with immunoblot which was separated in BRCA1 knockdown DU145 cells with siBRCA1-1 after treatment with CPT (20 μM) for 1 h and recovery for indicated time. (C) Co-IP was performed to detect interaction between PTEN and BRCA1 in 293T^senp−/−^ cells which were overexpressed indicated plasmids for 48 h and treated with CPT (20 μM). (D-E) SIM location in BRCA1 were predicted with JASSA and GPS-SUMO software. We mainly focused on those SIMs marked with red font which were located in exon11 of BRCA1. (F) Schematic representation of mutation of selected SIMs. All those amino acids of predicted SIMs were mutated into alanine, except SIM5 (5-1 and 5-2) which had been deleted. Relative location of predicted SIMs in BRCA1 protein were shown at lower panel. **Figure 5-figure supplement 1-source data 1. Western blots of Figure 5-figure supplement 1.**

**Figure 6-figure supplement 1.**
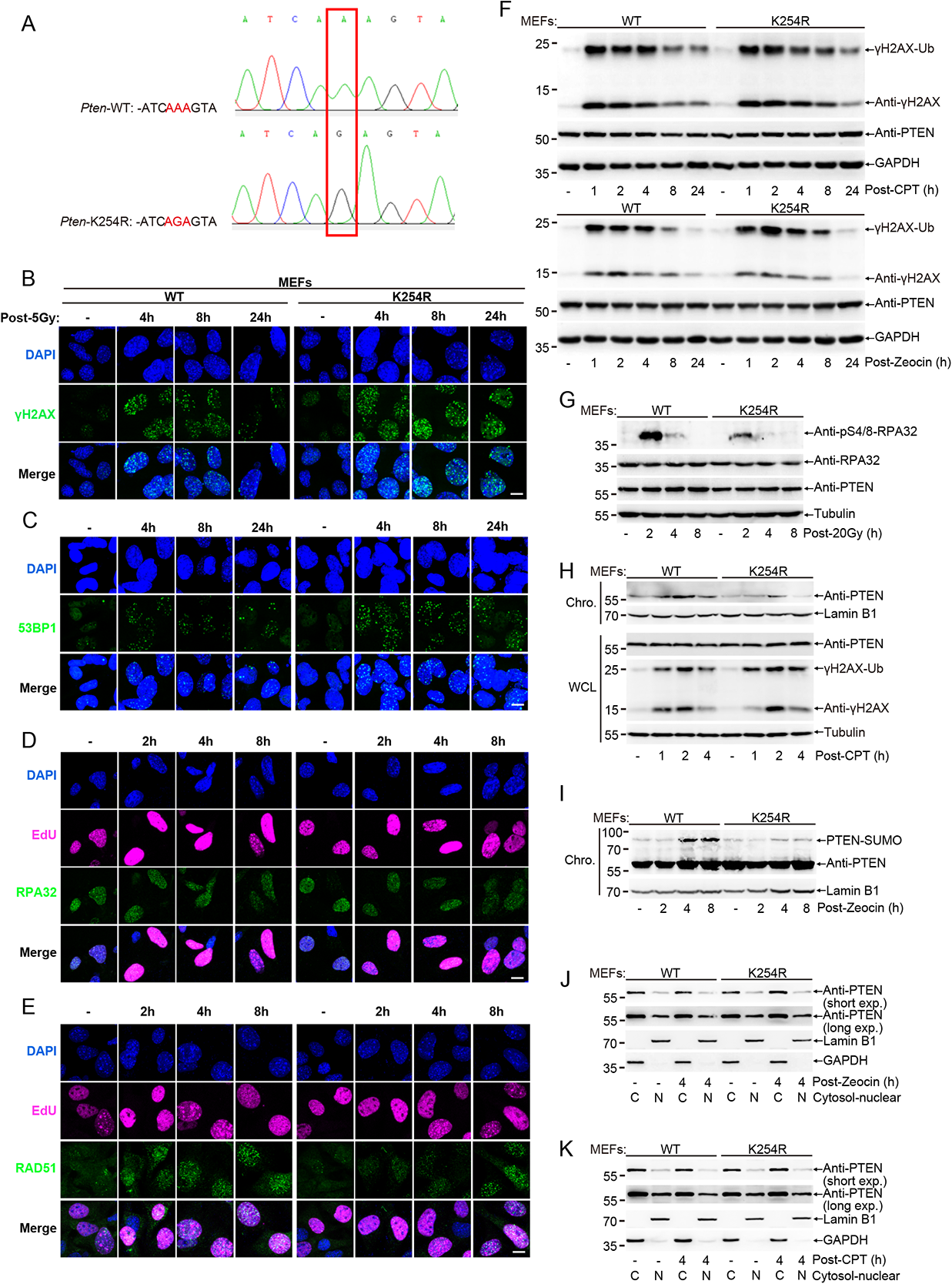
HR repair is impaired by SUMO-deficient PTEN *in vivo*. (A) Sequencing of *Pten*^WT^ and *Pten*^K254R^ knockin mouse. (B-E) Representative images of 53BP1, γH2AX, RAD51 and RPA32 foci of MEFs corresponding to Fig. 6A-D which were treated with 5 Gy and recovery for indicated time. (F) Immunoblot of γH2AX from MEFs treated with CPT (20 μM) or Zeocin (200 μg/mL) for 1 h and recovery for indicated time. (G) Immunoblot of pS4/8-RPA32 from MEFs treated with 20 Gy and recovery for indicated time. (H) Immunoblot of chromatin loaded PTEN which was separated from MEFs treated with CPT (20 μM) for 1 h and recovery for indicated time. (I) Immunoblot of chromatin loaded PTEN which was separated from MEFs treated with Zeocin (400 μg/mL) for 1 h and recovery for indicated time under harsh condition. (J, K) Nuclear-cytosol separation of MEFs post Zeocin (200 μg/mL) and CPT(20 μM) for 1 h and recovery for 4 h and then PTEN was detected with immunoblot. **Figure 6-figure supplement 1-source data 1. Western blots of Figure 6-figure supplement 1.**

**Figure 7-figure supplement 1.**
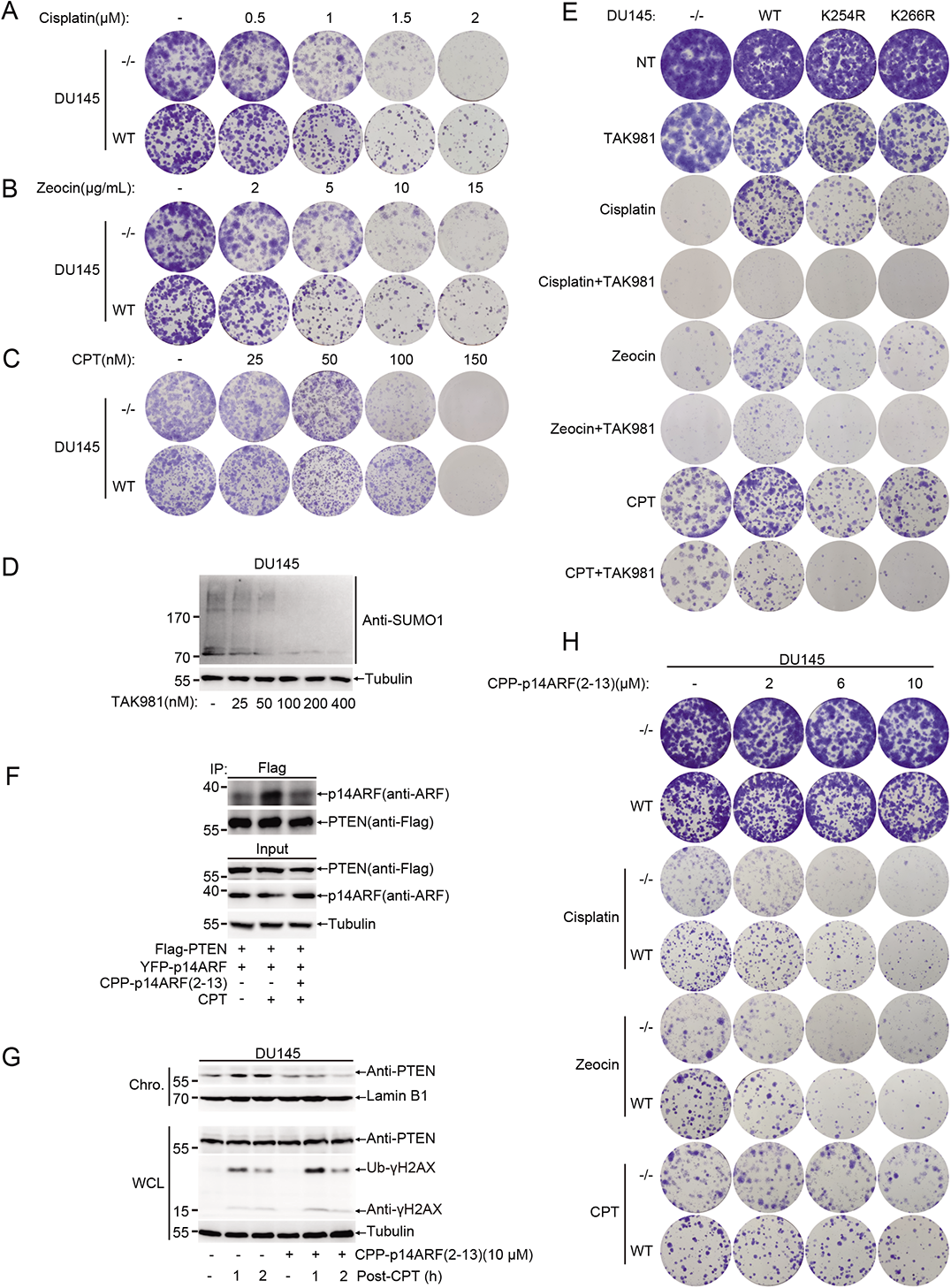
Blocking PTEN SUMOylation pathway sensitizes tumor cells to DNA damage reagents. (A-C) Representative colony images of DU145-PTEN^−/−^ and PTEN^WT^ cells treated with or without different doses of Cisplatin, Zeocin or CPT corresponding to Fig. 7A-C. (D) Immunoblot of total SUMO1 conjugates in DU145 cells treated with different doses of TAK981. (E) Representative colony images corresponding to Fig. 7D-G. (F) 293T cells were transfected with Flag-PTEN and YFP-p14ARF for 48 h and then treated with or without CPP-p14ARF(2-13) for 4 h. CPT (20 μM) was then added into culture medium for 1 h. Cells were collected after another 2 h. Interaction between PTEN and p14ARF were identified with Co-IP. (G) DU145-PTEN^WT^ cells were treated with or without CPP-p14ARF(2-13) for 4 h and then cells were treated with CPT (20 μM) for 1 h and recovery indicated time. Chromatin loaded PTEN was separated and detected with immunoblot. (H) Representative colony images corresponding to Fig.7I-L. 500 cells were seeded in 12-well plate for all colony assays except (C) in which 1000 cells were seeded at the beginning. **Figure 7-figure supplement 1-source data 1. Western blots of Figure 7-figure supplement 1.**

