## Supplementary file 1 for "SUMOylation of PTEN promotes DNA end resection through directly dephosphorylating 53BP1 in homologous recombination repair"

**Supplementary file 1A. List of antibodies used in this study.**

| Antibodies | | | | |
|---|---|---|---|---|
| **Name** | **Dilution Ratio** | **Application** | **Catalog Number** | **Manufacture** |
| anti-PTEN | 1:50 | IF | ab79156 | Abcam |
| anti-PTEN | 1:1000 | WB | #9559 | CST |
| anti-RAD51 | 1:200, 1:1000 | IF,WB | ab176458 | Abcam |
| anti-RPA32 | 1:200, 1:1000 | IF,WB | ab2175 | Abcam |
| anti-RPA32 | 1:200, 1:1000 | IF,WB | ab76420 | Abcam |
| anti-pS4/8-RPA32 | 1:1000 | IF | A300-245A-M | Bethyl |
| anti-BRCA1 | 1:200 | WB | sc-6954 | Santa Cruz Biotechnology |
| anti-53BP1 | 1:400,1:1000 | IF,WB | NB100-304 | Novus Biologicals |
| anti-pT543-53BP1 | 1:400,1:1000 | IF,WB | #3428 | CST |
| anti-pS25/29-53BP1 | 1:1000 | WB | #2674 | CST |
| anti-pS25-53BP1 | 1:200 | IF | NB100-1803 | Novus Biologicals |
| anti-p14ARF | 1:1000 | WB | A300-340A | Bethyl |
| anti-γH2AX | 1:200, 1:1000 | IF,WB | #9718 | CST |
| anti-γH2AX | 1:200, 1:1000 | IF,WB | ab26350 | Abcam |
| anti-RIF1 | 1:200 | IF | A300-567A | Bethyl |
| anti-Cyclin A2 | 1:200 | IF | ab181591 | Abcam |
| anti-Cyclin A2 | 1:200 | IF | #MA1-180 | Thermo Fisher |
| anti-SUMO1 | 1:1000 | WB | ab32058 | Abcam |
| anti-GST | 1:5000 | WB | #CW0084 | CWbioTech |
| anti-GAPDH | 1:5000 | WB | #60004-1-Ig | ProteinTech |
| anti-Tubulin | 1:5000 | WB | #66031-1-Ig | ProteinTech |
| anti-Lamin B1 | 1:2000 | WB | #66095-1-Ig | ProteinTech |
| anti-His | 1:1000 | WB | #66005-1-Ig | ProteinTech |
| anti-HA | 1:1000 | WB | #A448-101L | Covance |
| anti-Myc | 1:1000 | WB | #2276 | CST |
| anti-Flag | 1:1000 | WB | F1804 | Sigma |
| Alexa Fluor 488 Goat anti-Rabbit IgG | 1:1000 | IF | A27034 | Invitrogen |
| Alexa Fluor 568 Goat anti-mouse IgG | 1:1000 | IF | A-11004 | Invitrogen |

| | | | | |
|---|---|---|---|---|
| Alexa Fluor 488 Goat anti-mouse IgG | 1:1000 | IF | A28175 | Invitrogen |
| Alexa Fluor 568 Goat anti-Rabbit IgG | 1:1000 | IF | A-11011 | Invitrogen |
| normal mouse IgG | 1:250 | IP | sc-2025 | Santa Cruz Biotechnology |
| normal rabbit IgG | 1:250 | IP | sc-2027 | Santa Cruz Biotechnology |

### Supplementary file 1B. Sequences of shRNA, siRNA and sgRNA used in this study.

| **Sequences of shRNA, siRNA and sgRNA** | | |
|---|---|---|
| **Name** | **Sequences** | |
| | **sense（5'-3'）** | **antisense（5'-3'）** |
| shBRCA1-1 | GGAGCTCATTAAGGTTGTTGA | GGAGCTCATTAAGGTTGTTGA |
| siBRCA1-1 | CAGCUACCCUUCCAUCAUA | TATGATGGAAGGGTAGCTG |
| siControl | CAGCAGTTTATTACTCACTAA | TTAGTGAGTAATAAACTGCTG |
| shPTEN | CCACAAATGAAGGGATATAAA | CCACAAATGAAGGGATATAAA |
| shp14ARF-1 | GAAGACCAGGTCATGATGAT | GAAGACCAGGTCATGATGAT |
| shp14ARF-2 | GAACATGGTGCGCAGGTTCT | GAACATGGTGCGCAGGTTCT |
| sgPTEN-1 | CACCGTTATCCAAACATTATTGC TA | AAACTAGCAATAATGTTTGGATA AC |
| sgPTEN-2 | CACCGGCTAACGATCTCTTTGA TGA | AAACTCATCAAAGAGATCGTTAG C C |
